# Grapevine microbiota reflect diversity among compartments and complex interactions within and among root and shoot systems

**DOI:** 10.1101/2020.11.02.365197

**Authors:** Joel F. Swift, Megan E. Hall, Zachary N. Harris, Misha T. Kwasniewski, Allison J. Miller

**Affiliations:** Saint Louis University, Department of Biology, 3507 Laclede Ave, St. Louis, MO 63103, USA; Donald Danforth Plant Science Center, 975 North Warson Road, St. Louis, MO 63132, USA; University of Missouri, Division of Plant Sciences, Agriculture Bldg, 52, Columbia, MO 65201, USA; The Pennsylvania State University, College of Agricultural Sciences Department of Food Science, 326 Rodney A. Erickson Food Science Building, University Park, PA 16802, USA

**Keywords:** Grafting, Grapevines, Rootstock, Sour rot, Plant compartments, Bacteria, Fungi

## Abstract

**Background:** Within an individual plant, different compartments (e.g. roots, leaves, fruits) host distinct communities of microorganisms due to variation in structural characteristics and resource availability. Grafting, which joins the root system of one individual with the shoot system of a second genetically distinct individual, has the potential to bring the microbial communities of different genotypes together. An important question is the extent to which unique root system and shoot system genotypes, when grafted together, influence the microbiota of the graft partner. Our study sought to answer this question by utilizing an experimental vineyard composed of ‘Chambourcin’ vines growing ungrafted and grafted to three different rootstocks, replicated across three irrigation treatments. We characterized bacterial and fungal communities in roots, leaves, and berries, as well as surrounding soil. Our objectives were to (1) characterize the microbiota of compartments within the root system (roots and adjacent soil) and the shoot system (leaves and berries), (2) determine the influence of rootstock genotypes, irrigation, and their interaction on the microbiota of aboveground and belowground compartments, and (3) investigate the distribution of microorganisms implicated in the late-season grapevine bunch rot disease sour rot (*Acetobacterales* and *Saccharomycetes*).

**Results:** Compartments were significantly differentiated in bacterial and fungal richness and composition. Abundance-based machine learning accurately predicted the compartment and differential abundance analysis showed a large portion of taxa differed significantly across compartments. Rootstock genotypes did not differ significantly in microbial community richness or composition; however, individual microbial taxa exhibited significant differences in abundance based on rootstock and irrigation treatment. The relative abundance of *Acetobacterales* and *Saccharomycetes* in the berry was influenced by complex interactions among rootstock genotype and irrigation.

**Conclusion:** Our results indicate that grapevine compartments retain distinct core microbiota regardless of the rootstock to which they are grafted. While rootstock genotype generally had a subtle impact on global patterns of microbial diversity, we found associations between rootstock genotypes and specific groups of microorganisms. Further experimental validation is needed in order to understand how associations with these microorganisms impacts a vine’s susceptibility to sour rot upon damage and whether the characteristics of wine are impacted.

## Background

As plants have evolved over millennia, so too have their biological interactions with microorganisms, including bacteria, fungi, archaea, and viruses. Plants have multiple compartments (e.g. roots, leaves, fruits, etc.), each of which offer unique habitats for microorganisms. Plant compartments differ in structural characteristics, micro-environmental conditions, and resource availability which differentially regulate their microbiota (Chi *et al*. 2005; Vorholt 2012; Bulgarelli *et al*. 2013; Martins *et al*. 2013; Turner *et al*. 2013; Hacquard *et al*. 2015). The microbiota of plant compartments contribute to many essential processes, including nutrient acquisition (Bulgarelli *et al*. 2013) and adaptation to novel soil conditions (Lau and Lennon 2012; Keymer and Lankau 2017), among others. The composition of microbiota within individual plant compartments reflects dynamic interactions of plant genotype, development, and local environment; however, the extent to which the microbiota in one compartment (e.g., the root) shapes the microbiota of other parts of the plant (e.g., leaves or fruit) is not well known.

Many factors influence the composition and diversity of plant microbiota, including geographic location of the plant, host plant genotype, and biotic and abiotic stresses. Biogeography is a predominant influence on plant microbiota (Bokulich *et al*. 2014; Coleman-Derr *et al*. 2016; Walters *et al*. 2018), reflecting unique soil microorganism communities across space (Green *et al*. 2004; Fierer and Jackson 2006). Plant host genotype also shapes plant microbiota (Peiffer *et al*. 2013; Mahoney *et al*. 2017; Jiang *et al*. 2017; Fitzpatrick *et al*. 2018). For example, within four days of planting aseptically cleaned seeds of nine cotton (*Gossypium hirsutum*) cultivars, Adams and Kloepper (2002) found differences in bacterial endophyte composition between cultivars. Genotype-specific microbiota have been identified in numerous other genera including *Helianthus* (Leff *et al*. 2016), *Solanum* (Inceoǧlu *et al*. 2010), *Triticum* (Mahoney *et al*. 2017), *Vaccinium* (Jiang *et al*. 2017), and *Zea* (Bouffaud *et al*. 2012; Peiffer *et al*. 2013; Szoboszlay *et al*. 2015). The mechanism commonly proposed to explain the observed differences by genotype is altered root exudation profiles (Micallef *et al*. 2009; Sasse *et al*. 2018; Walters *et al*. 2018). Plants are also capable of modifying their exudation patterns in response to biotic and abiotic stresses, thereby recruiting microorganisms that can lessen damage to plant health (Naylor and Coleman-Derr 2018; Rolfe *et al*. 2019). Both drought and pathogen infection cause strong and lasting effects on root exudate profiles that not only impact the individual plant, but also subsequent generations grown in the same location (Gargallo-Garriga *et al*. 2018; Yuan *et al*. 2018).

Most studies investigating the factors influencing the microbiome of crops are focused on annual species; however, woody perennials are economically important worldwide (Food and Agriculture Organization 2016) and present their own intricacies (Warschefsky *et al*. 2016). For example, bacterial richness and composition of root-associated microbiota have been shown to shift over the lifetime of perennial plants (Wagner *et al*. 2016). Additionally, many woody perennial species are grafted, a horticultural practice that joins the root system (rootstock) of one individual to the shoot system (scion) of another (Pina and Errea 2005; Mudge *et al*. 2009). Grafting physically connects individuals with distinct genomes, allowing genome to genome interactions (Gaut *et al*. 2019). Given that plant genotype influences the microbiome, an important question is the extent to which unique genotypes, when grafted together, influence the microbiome of the graft partner.

Grafted grapevines present an ideal model to understand interactions among root and shoot systems and their effects on the microbiota in different plant compartments. The European grapevine (*Vitis vinifera*) is one of the most economically important berry crop species in the world (Food and Agriculture Organization 2016). Since the spread of the root-destroying Phylloxera aphid from North America to Europe in the 1800s (Kliman 2010), grafting to phylloxera-resistant rootstocks has become ubiquitous in the growing of grapevines. Today more than 80% of vineyards employ grafting (Ollat *et al*. 2016). Previous research on grafted grapevines has shown that rootstock genotypes influence scion phenotypes such as leaf ionomic profiles (Lecourt *et al*. 2015; Gautier *et al*. 2020), shoot gene expression (Cookson and Ollat 2013), and leaf morphology (Migicovsky *et al*. 2019). Interestingly, grapevine rootstock genotypes have distinct bacterial rhizosphere communities (D’Amico et al. 2018; Marasco et al. 2018), and the influence of rootstock genotype on root-associated microbiota appears to increase with vine age (Berlanas *et al*. 2019). These studies suggest that rootstock genotypes grafted to a common scion retain the ability to recruit distinct root-associated microbiota and this pattern becomes stronger across the lifespan of the grapevine. The extent to which rootstock genotype influences the microbiota of scion compartments (e.g. leaves and berries) is an open question that requires sampling and sequencing from both above- and below-ground compartments of grafted plants.

Many grapevine rootstocks have been bred primarily for resistance to specific soil-borne pests and pathogens (Reisch *et al*. 2012); however, some microorganisms of concern for vine health are associated primarily with the shoot system (Wilcox *et al*. 2015). For example, sour rot is a late-season bunch rot disease that causes loss of production across many growing regions, particularly those with high amounts of summer rainfall (Wilcox *et al*. 2015). Sour rot is typified by oxidation of grape berry skin and loss of berry integrity prior to harvest and is accompanied by a strong odor of acetic acid and the presence of *Drosophila* spp. (Bisiach *et al*. 1982; Ioriatti *et al*. 2018). Berry clusters affected by sour rot result in altered fermentation, changes in wine characteristics, and generally lower wine quality (Zoecklein *et al*. 2000; Barata, Campo, *et al*. 2011; Barata, Pais, *et al*. 2011). This disease has been linked to a four way interaction between a susceptible grapevine host, *Drosophila* fruit flies, acetic acid bacteria (e.g. *Acetobacter* spp. and *Gluconobacter* spp.) and various yeast species (*Hanseniaspora uvarum, Candida* spp., and *Pichia* spp.; Hall *et al*. 2018). The bacteria and yeast associated with sour rot are commonly found as endo- and epiphytes of asymptomatic berry clusters (Pinto *et al*. 2014; Hall and Wilcox 2019); however, as symptoms of sour rot develop the microbiota of infected clusters the typically show an elevated abundance of acetic acid bacteria, particularly *Acetobacter* spp. (Hall *et al*. 2019). In addition to being causative agents in sour rot, acetic acid bacteria and non-*Saccharomyces* yeast species are important to the winemaking process. *Acetobacter* spp. and *Gluconobacter* spp. are able to both oxidize sugars into acetic acid and contribute to higher volatile acidity levels (Barata, Campo, *et al*. 2011; A. Barata *et al*. 2012). While *Hanseniaspora uvarum, Candida* spp., and *Pichia* spp. are considered spoilage yeasts in winemaking via production of ethyl acetate and film formation of stored wine (A. Barata *et al*. 2012; Malfeito-Ferreira 2019) and can dominate spontaneous fermentation products (Portillo and Mas 2016). The role of rootstocks in shaping disease-causing bacterial and yeast populations in the shoot system is not well-known.

This study aims to advance understanding of the influence of rootstock genotype on microbiota of grafted grapevines. Samples were collected from an experimental vineyard composed of the grapevine cultivar ‘Chambourcin’ as the scion, grown ungrafted and grafted to three different rootstocks. We surveyed multiple replicates of each ‘Chambourcin’ scion/rootstock combination, and collected samples from four distinct compartments for each vine: leaves, berries, root tissue (with rhizosphere attached) and soil. We quantified the bacterial and fungal members of each sample using amplicon sequencing. Our objectives were to (1) characterize the microbiota of both the root and shoot system, (2) determine the association of rootstock genotype, irrigation treatment, and their interaction on the microbiota of above and belowground compartments, and (3) investigate the distribution of microorganism groups implicated in the late-season grapevine bunch rot disease sour rot (*Acetobacterales* and *Saccharomycetes*)

## Materials and methods

### Experimental design and sample collection

The rootstock experimental vineyard was established in 2009 at the University of Missouri Southwest Research Center Agricultural Experimental Station in Mount Vernon, Missouri USA (37.074, -93.879). The ∼8000 m^2^ vineyard is planted with vine rows running from east to west, and experiences mean annual rainfall of 1066.8 mm, mean annual temperature of 15.6°C, and mean annual growing degree days of 461 (Maimaitiyiming *et al*. 2017; https://gwi.missouri.edu/IPMreports/2019-no8_GrowingDegree.htm). Soil is a combination of sandy loam, silt loam, and loam with an average pH of 7 (Maimaitiyiming *et al*. 2017; Table S1). A subplot within the vineyard consists of the *Vitis* interspecific hybrid, ‘Chambourcin’, growing ungrafted and grafted to three different rootstocks: 1103 Paulson (‘1103P’; *Vitis berlandieri* × *V. rupestris*), 3309 Courdec (‘3309C’; *V. riparia* × *V. rupestris*), and Selection Oppenheim 4 (‘SO4’; *V. berlandieri* × *V. riparia*). Each scion/rootstock combination is replicated 72 times in a randomized block experimental design (Figure 1). Two additional vines are planted at the beginning, end, and middle point of a row to buffer experimental vines against edge effects and irrigation risers. One of three different irrigation treatments are applied by row in a randomized order 1) no irrigation, 2) full replacement of evapotranspiration as calculated on a rolling weekly basis, and 3) half replacement of evapotranspiration (Figure 1). This vineyard undergoes chemical spray applications for fungal diseases (e.g. downy mildew, black rot, Botrytis bunch rot, and Phomopsis cane and leaf spot), consistent with industry practices. Without fungicide applications, vines become infected with fungal pathogens, resulting in loss of fruit, leaves, and potential death of the vines.

**Figure 1.**
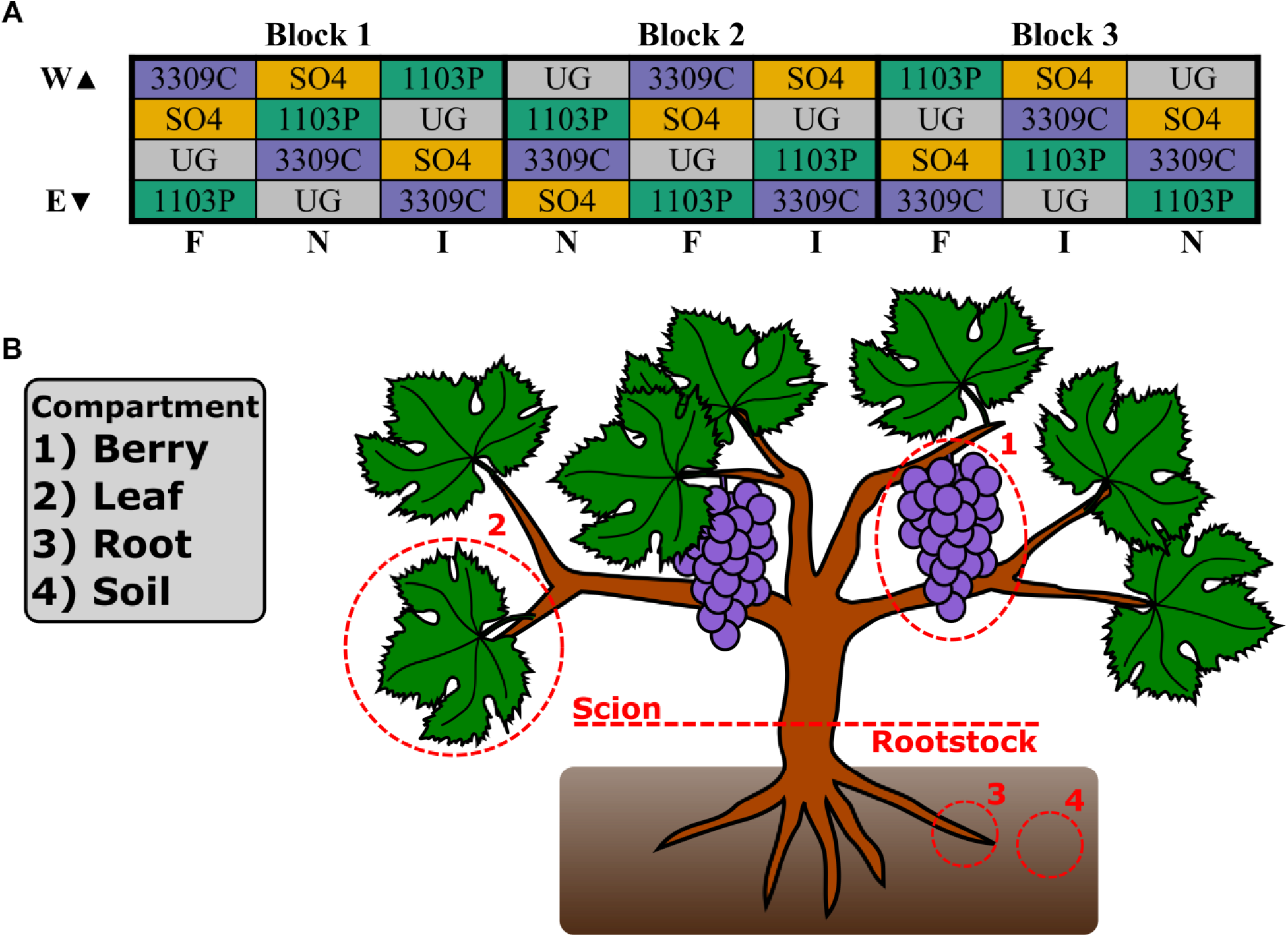
Experimental design. A) Vineyard layout at the University of Missouri Southwest Research Center in Mount Vernon, MO consisting of 288 vines grafted to one of three rootstocks (‘1103P’, 1103 Paulson; ‘3309C’, 3309 Courdec; ‘SO4’, Selection Oppenheim 4) or ungrafted (UG). Each colored cell represents four vines; irrigation treatments (F, Full replacement of evapotranspiration; I, reduced replacement; N, unirrigated) and experimental blocks are listed along the bottom and top of the grid, respectively. B) Depiction of a grafted grapevine with bulk soil and different compartments highlighted, numbers correspond to legend (left).

Samples were collected from 72 vines (four scion/rootstock combinations × three irrigation treatments × six replicates per scion/rootstock/irrigation combination) on September 18, 2018, coinciding with the timing of commercial harvest (E-L #38: Jones 2003). For each vine, samples were collected from bulk soil, rhizosphere, leaves, and one berry cluster (Figure 1). Bulk soil was collected from at least 3 cm below ground level with a shovel or hand trowel at the base of the stem, and passed through a sieve (American Standard No. 16; 1.18 mm pore size). Roots were collected at a depth of 20-30 cm by manually removing roots from the soil. Five leaves 8-12 cm in diameter were collected at roughly the same position along the shoot and height on the vine. Berries were collected as an intact cluster, free of damage or apparent disease symptoms. All collections were made using sterilized equipment and conditions. For each treatment (Figure 1; colored cells), samples were collected from the two inner of a set of four plants. In block three, samples were collected on an individual vine basis, whereas in blocks one and two samples were homogenized per treatment (rootstock genotype) in the field. In total, 192 samples (48 from each compartment type) were collected in polyethylene bags, placed in a cooler with dry ice for transport, and stored at -80°C until DNA extraction.

### DNA extraction, amplification, and sequencing

DNA extractions for all samples were performed using the DNeasy PowerSoil Kit (Qiagen) following the manufacturer’s protocol with two modifications; plant tissue samples had 150 mg per extraction and a 10-minute (70°C) incubation step prior to homogenization with a bead mill (Retsch MM 400). For berry cluster samples, two berries of equal size were placed into a sterilized stainless-steel grinding jar with a stainless-steel ball (Retsch), frozen in liquid nitrogen, and then pulverized prior to homogenization. Leaf and root samples were finely chopped with a sterile scalpel prior to homogenization. All extracts were quantified using a DS-11 spectrophotometer (Denovix).

Library preparation and amplicon sequencing was conducted at the Environmental Sample Preparation and Sequencing Facility at Argonne National Laboratory. The V4 region of the 16S rRNA gene (515F-806R; Parada *et al*. 2016 and Apprill *et al*. 2015, respectively) and the internal transcribed region (ITS1f & ITS2; Smith and Peay 2014) were separately amplified with region-specific primers that include sequencer adapter sequences used in the Illumina flowcell (Caporaso *et al*. 2011, 2012). The forward amplification primer contained a twelve base barcode sequence to allow pooling of samples onto a single sequencing lane (Caporaso *et al*. 2011, 2012). To each 96-well plate two negative controls (extraction and PCR) and one positive control, ZymoBIOMICS™ Microbial community DNA standard (Cat No. D6305) were added. Each 25 µL PCR reaction contained 9.5 µL of MO BIO PCR Water, 12.5 µL of QuantaBio’s AccuStart II PCR ToughMix (2x concentration, 1x final), 1 µL barcode tagged Forward Primer (5 µM concentration, 200 pM final), 1 µL Reverse Primer (5 µM concentration, 200 pM final), and 1 µL of template DNA. For 16S rRNA gene PCRs peptide-nucleic-acid blockers are used to reduce non-target amplicons originating from chloroplast and mitochondria (Lundberg *et al*. 2013), 1 µL (5 µM concentration, 200 pM final) of each is used. The conditions for the 16S rRNA gene PCR are as follows: 94°C for 3 minutes, with 35 cycles at 94°C for 45s, 50°C for 60s, and 72°C for 90s; with a final extension of 10 min at 72°C. The conditions for the ITS PCR are as follows: 94°C for 1 minute, with 35 cycles at 94°C for 30s, 52°C for 30s, and 68°C for 30s; with a final extension of 10 min at 68°C. Amplicons are then quantified using PicoGreen (Invitrogen) and a plate reader (Infinite 200 PRO, Tecan). Once quantified, volumes of each of the products are pooled in equimolar amounts. This pool was then cleaned using AMPure XP Beads (Beckman Coulter), and quantified using a fluorometer (Qubit, Invitrogen). After quantification, the molarity of the pool was determined and diluted down to 2 nM, denatured, and then diluted to a final concentration of 6.75 pM with a 10% PhiX spike. Sequencing was conducted on an Illumina MiSeq, 2×151bp PE for 16S rRNA gene and 2×250b PE for ITS using customized sequencing primers and procedures (Caporaso *et al*. 2012).

### Bioinformatic processing

Data processing and preliminary analyses were conducted in QIIME2 (Bolyen *et al*. 2018). Samples were demultiplexed (*qiime demux emp-paired*) according to barcode sequence. For 16S rRNA gene sequences, the QIIME2 plugin DADA2 (Callahan *et al*. 2016) was used to denoise, dereplicate, and filter chimeric sequences. The first 13 nucleotides (nt) of each sequence (forward and reverse) was trimmed and truncated at 150 nt to remove lower quality bases. For ITS sequences, Cutadapt (Martin 2011) was used to remove primer sequences prior to using the DADA2 plugin. The first 12 nt of the 3′ end of each sequence were trimmed, the 5′ end of sequences were not truncated to preserve biologically relevant length variation (Schoch *et al*. 2012), and a max expected error rate of 3 was used. The result of DADA2 processing was a table of Amplicon Sequence Variants (ASV). Workflows using ASVs perform similar to OTU based methods with the benefits of comparability across studies (using the same primers) and increased taxonomic resolution (Callahan *et al*. 2017; Glassman and Martiny 2018). Taxonomic classification of ASVs was conducted with a naive Bayes classifier trained on either the SILVA (v.132; trimmed to the V4 region; Yilmaz *et al*. 2014) or UNITE database (v.8.0; Unite Community 2018) for bacteria and fungi, respectively. Bacterial and fungal ASVs not assigned to a phylum were removed along with bacterial ASVs assigned to mitochondria or chloroplasts. ASVs with less than 0.1% of the total filtered reads, which tend to inflate diversity metrics (Bokulich *et al*. 2013), were removed. Negative and positive control samples were processed in a similar manner but we did not remove bacterial ASVs assigned to mitochondria or chloroplasts or filter those at less than 0.1% of filtered reads (Figure S1).

### Statistical analyses

Samples were rarefied to 1,500 and 5,000 sequences for bacteria and fungi, respectively. Alpha diversity and beta diversity metrics were calculated in phyloseq (McMurdie and Holmes 2013). For bacterial samples, we calculated three alpha diversity metrics (inverse Simpson’s, Shannon’s, and Faith’s phylogenetic diversity indices) and the beta diversity metric UniFrac (both weighted and unweighted), which considers phylogenetic relatedness between samples in calculations (Lozupone and Knight 2005). For fungal samples, we calculated inverse Simpson’s and Shannon’s diversity indices and Bray-Curtis dissimilarity, a beta-diversity metric. Principal coordinates analysis was used to visualize sample relationships. Venn diagrams were generated by determining the intersection of ASV lists per compartment (URL: http://bioinformatics.psb.ugent.be/webtools/Venn/).

Statistics were calculated in base R (v.3.6.1; R Core Team 2019) and in various R packages. Figures were generated with ggplot2 (v.3.2.1; Wickham 2016) and arranged using ggpubr (https://github.com/kassambara/ggpubr/). In order to test for significant differences between alpha diversity means by compartment, a Tukey’s honestly significant difference (HSD) test was applied. Linear models and PERMANOVA tests were run using the formula: response variable ∼ Rootstock genotype × Compartment × Irrigation + Block. For each alpha diversity metric, a linear model was fit using lm and assessed using type-III ANOVA with the car package (v3.0-3; Fox and Weisberg 2019). Beta diversity metrics were subjected to PERMANOVA tests with adonis in the vegan package (Anderson, M. J. 2001; Oksanen *et al*. 2019) using 10,000 permutations.

In order to investigate the distribution of bacterial and fungal taxa associated with the late-season bunch rot disease sour rot, we used the *subset_taxa* function in phyloseq to extract ASVs from the rarified datasets that assigned to the bacterial family *Acetobacterales* or the fungal class *Saccharomycetes*. Relative abundance of these taxa across compartments, and for rootstock and irrigation treatments, were visualized using boxplots. We ran linear models using the full experimental design formula (Rootstock genotype × Compartment × Irrigation + Block) with the abundance of *Acetobacterales* or *Saccharomycetes* as the response variable. We conducted post-hoc comparisons of means using emmeans (Lenth *et al*. 2020), correcting for multiple comparisons, to understand which comparisons of significant factors were driving the abundance of these taxa. To test for correlation between the relative abundance of *Acetobacterales* and *Saccharomycetes*, we conducted a spearman rank based correlation using the *stat_cor* function from the ggpubr package (https://rpkgs.datanovia.com/ggpubr/).

In order to determine whether each rootstock had a predictable impact on the microbiota we used a two-pronged approach: 1) machine learning to identify patterns across the dataset; and 2) differential abundance analysis to identify individual microorganisms. First, for machine learning we used ranger’s implementation of random forest (v.0.11.2; Wright and Ziegler 2017) and tuned hyperparameters (number of trees, minimum node size, and number of features available at each node) with Caret (v.6.0-84; Kuhn 2008) on 80% of the dataset (20% withheld for testing). The optimal hyperparameters were selected by iteratively assessing the performance on out-of-bag samples for a given parameter set. We then used this final model to predict the label, either rootstock or compartment or both, on the withheld testing data assessing the prediction accuracy. Data were visualized using tile plots from the output confusion matrix. Second, for differential abundance analysis, we used unrarefied reads and removed ASVs that were not represented by a depth of least 25 reads in more than 10% of the samples to conduct differential abundance modeling with DESeq2 (v.1.24.0; Love *et al*. 2014). DESeq2 was fit using the following model: response variable ∼ Rootstock genotype (R) + Compartment (C) + Irrigation (I) + R × C + I × R + I × C + Block (B). For each factor we extracted the number of ASVs that showed a significant pattern of differential abundance as well as their fold changes (Log_2_) to generate summary plots.

## Results

We generated bacterial (16s rRNA) and fungal (ITS) amplicon sequence data for bulk soil, roots, leaves and berries of 72 grapevines. For bacteria, this study produced 13,003,903 reads, of which 10,647,004 reads remained following quality and chimera filtering. Reads collapsed into 20,602 bacterial ASVs. In order to remove host contamination, bacterial ASVs assigned to mitochondria and chloroplasts were removed (9.1% and 31.4% of reads, respectively). ASVs with less than 0.1% mean sample depth were removed, resulting in 8,199 ASVs and 6,155,632 reads for bacteria. We used rarefaction to normalize the number of reads per sample, 1,500 reads per sample, resulting in the removal of seven samples (Figure S2A). For fungi, we generated 11,434,967 reads; 5,748,318 reads passed quality and chimera filtering, and these collapsed into 2,475 ASVs. After mean sample depth filtering the resulting fungal dataset included 1,429 ASVs comprised of 5,735,409 reads. Finally, we rarefied fungal samples at 5,000 reads, resulting in 11 samples being removed (Figure S2B).

### Bacterial and fungal community composition and richness strongly associate with plant compartment

Principal coordinates analyses for bacterial unweighted UniFrac distances and fungal Bray-Curtis distance show clear clusters by compartment, with the first axis separating above and belowground compartments, the second axis separating soil and roots, and the third axis begins to pull apart berries and leaves (Figure 2A and 2D; For axis three see Figure S5). For bacteria the first and second principal coordinate axes explained 32.5% and 5.4% and for fungi the first and second principal coordinate axes explained 44.8% and 8.0% (Figure 2A and 2D). PERMANOVA tests corroborated the PCoA results: for bacteria, compartment was the only significant factor (unweighted UniFrac: *F*_3,176_ = 40.7, *p*<0.001; Table 1) while fungi showed significant effects of compartment and some of its interactions (Table 1).

**Table 1.** Permutational Multivariate ANOVA results for bacterial and fungal communities using Unweighted UniFrac and Bray-Curtis distance metrics, respectively. Factors that are determined to be significant are bolded and interactions are denoted with the following abbrevations (R) rootstock, (C) compartment, and (I) irrigation.

| Factor | Unweighted UniFrac |  |  | Bray-Curtis |  |  |
| --- | --- | --- | --- | --- | --- | --- |
|  | Pseudo-*F* | *p*-value | R<sup>2</sup> | Pseudo-*F* | *p*-value | R<sup>2</sup> |
| Rootstock (R) | $F_{3,176} = 1.124$ | 0.252 | 0.011 | $F_{3,173} = \mathbf{2.178}$ | **0.013** | **0.016** |
| Compartment (C) | $F_{3,176} = \mathbf{40.720}$ | **< 0.001** | **0.407** | $F_{3,173} = \mathbf{74.320}$ | **< 0.001** | **0.550** |
| Irrigation (I) | $F_{2,177} = 1.107$ | 0.273 | 0.007 | $F_{2,174} = 1.203$ | 0.247 | 0.006 |
| Block | $F_{2,177} = 1.304$ | 0.147 | 0.009 | $F_{2,174} = 1.402$ | 0.151 | 0.007 |
| R × C | $F_{9,170} = 1.057$ | 0.326 | 0.032 | $F_{9,167} = \mathbf{1.459}$ | **0.040** | **0.032** |
| R × I | $F_{6,173} = 1.058$ | 0.324 | 0.021 | $F_{6,170} = 1.146$ | 0.261 | 0.017 |
| C × I | $F_{6,163} = 1.009$ | 0.417 | 0.020 | $F_{6,170} = 0.936$ | 0.526 | 0.014 |
| C × R × I | $F_{18,161} = 0.989$ | 0.501 | 0.059 | $F_{18,158} = 0.998$ | 0.473 | 0.044 |
| Residual |  |  | 0.433 |  |  | 0.313 |

**Figure 2.**
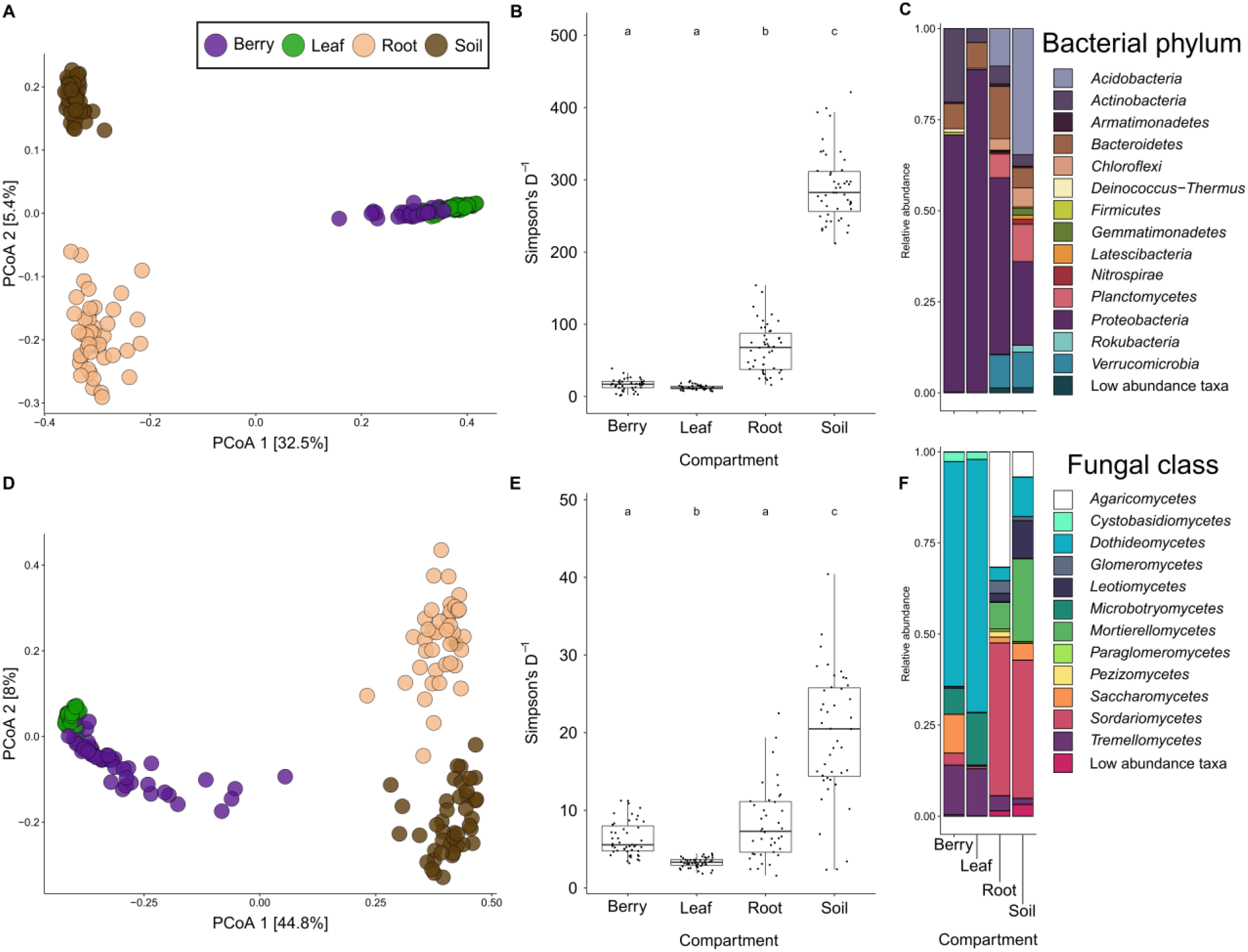
Inverse Simpson’s Diversity index of microorganism compartments for A) bacteria and D) fungal samples show a relative increase in diversity from above-ground to below-ground compartments. Principal coordinates analysis illustrates strong clustering of samples by compartment for B) bacterial (unweighted UniFrac: PERMANOVA *F*_3,176_ = 40.7, *p*<0.001) and E) fungal (Bray-Curtis: PERMANOVA *F*_3,176_ = 74.3, *p*<0.001). Taxonomic barplots for C) bacterial phyla and F) fungal classes reveals differences in microrganism community composition and structure by tissue.

Belowground compartments, soil and roots, were more diverse than aboveground compartments, leaves and berries, for both bacteria and fungi (Figure 2B and E). Bacterial communities were most diverse in soil, then roots, followed by berries and leaves, with mean inverse Simpson’s diversity values of 634.1, 437.9, 56.6, and 52.8, respectively (Figure 2B). Fungal communities showed similar patterns; however, fungal communities in berries were significantly more diverse than fungal communities in leaves with mean inverse Simpson’s diversity values 110.1, 66.0, 47.9, 29.1 for soil, roots, berries and leaves, respectively (Figure ED). For bacteria and fungi, linear models run on alpha diversity metrics demonstrated that compartment was significant across all metrics (p<0.001; Table S2 and S3). Venn diagrams showed a similar pattern to the alpha diversity metrics, while the intersections revealed that many ASVs are shared in multiple tissues and the soil for bacteria and fungi (Figure S4). Interestingly, while soil contained the highest number of ASVs, a sizable number of ASVs were found only in association with plant tissues and not in soil (∼26% and ∼32% for bacteria and fungi, respectively).

Differences in taxonomic profiles were apparent between compartments (Figure 2C, F). Bacterial samples of leaves and berries were dominated by Proteobacteria, with smaller proportions of Actinobacteria and Bacteroidetes (Figure 2C). Root and soil compartments were considerably more diverse but still showed a large proportion of the phylum Proteobacteria; additional phyla recovered in root and soil compartments included Acidobacteria (34.6% soil vs 10.3% root), Bacteroidetes, Verrucomicrobia, Planctomycetes, Actinobacteria, and Chloroflexi. Fungal samples of leaves and berries were also dominated by a few taxonomic classes, Dothideomycetes, Agaricomycetes, and Mortierellomycetes (Figure 2F). Berries showed the largest abundance of Saccharomycetes in comparison to other tissues (10.6% berries vs 4.5% soil). Root and soil compartments showed several additional fungal classes including Sordariomycetes, Tremellomycetes, Microbotrymycetes, Leotiomycetes, Cystobasidiomycetes, Glomeromycetes, Perizizomycetes, and Paraglomeromycetes.

### Rootstock genotype and irrigation have subtle effects on global community patterns

Global patterns of bacterial and fungal diversity, including alpha and beta diversity measured within compartments and across compartments within vines of specific scion/rootstock combinations, were generally similar regardless of rootstock (Figure 3). Alpha diversity metrics for bacteria showed no significant effect of rootstock genotype (Table S2). For fungi, inverse Simpson’s diversity index differed by rootstock (Table S3); post-hoc testing revealed that this variation was significantly described by rootstock when comparing ‘3309C’- and ‘SO4’-grafted vines in the soil samples (‘3309C’ vs ‘SO4’ soil: 10.369, t_127_ = 4.817, *P*<0.0001). Beta diversity analysis showed similar patterns to alpha diversity metrics (Table 1); for bacteria, rootstock and its interactions were non-significant. Fungi showed significant effects of rootstock and its interaction with compartment; however, rootstock and the interaction of rootstock and compartment showed much smaller impact than for compartment (Bray-Curtis: tissue *F*_3,173_ = 74.3, rootstock *F*_3,173_ = 2.2, compartment×rootstock *F*_9,167_ = 1.5; Table 1). Although irrigation treatments were imposed on the vines in this study (N; None, F; Full, and I; half replacement of evapotranspiration; Figure 1A), these did not have a large impact on the microorganism communities. Linear models fit to alpha and beta diversity metrics for both bacteria and fungi did not find irrigation to be a significant predictor (Table 1, S2, and S3).

**Figure 3.**
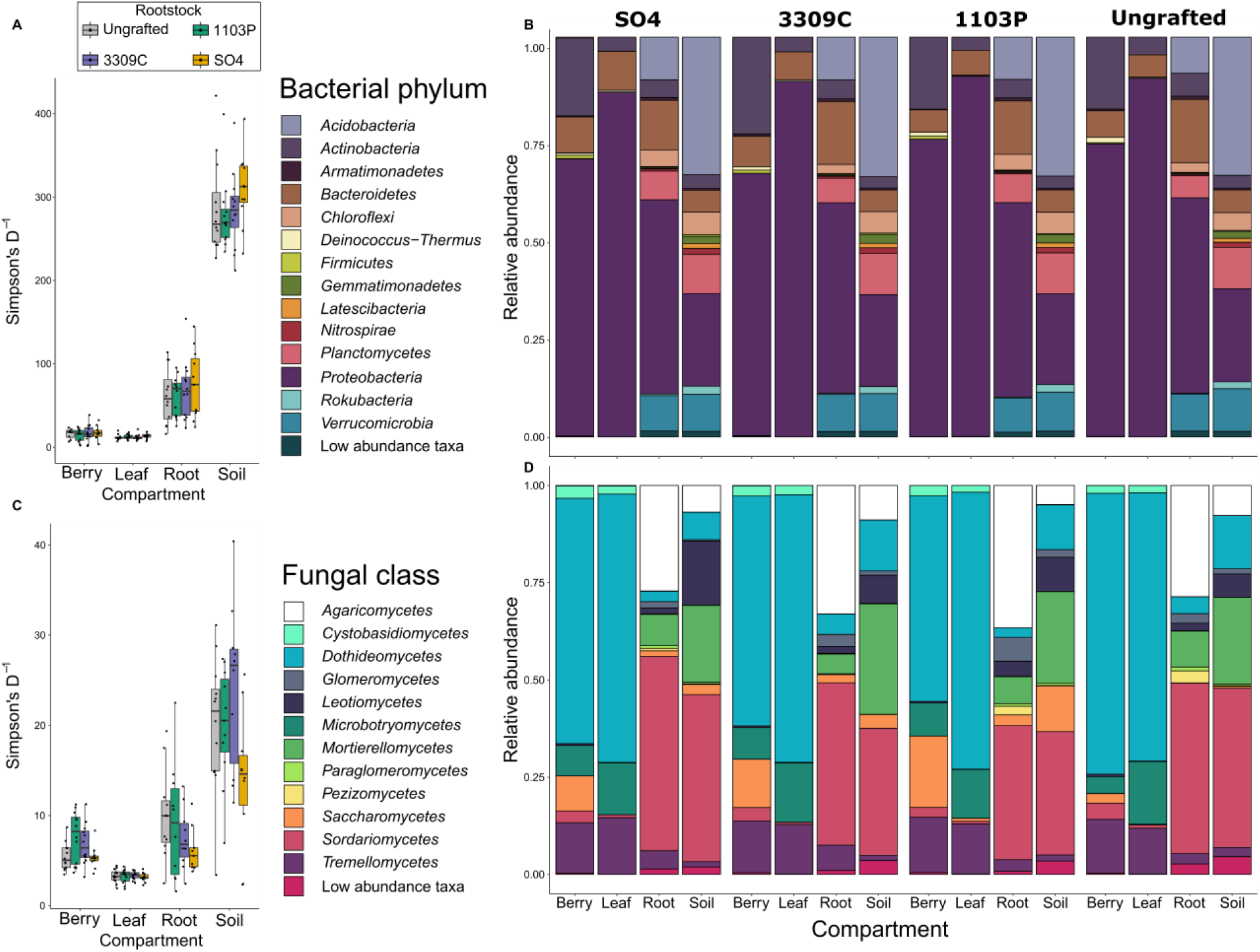
Inverse Simpson’s Diversity index and taxonomic barplots for microorganism compartments across rootstock genotypes for A-B) bacteria and C-D) fungal samples illustrate that rootstock has only subtle effects on global patterns of diversity and composition of microorganism communities of grapevines.

Taxonomic profiles were similar across rootstock genotypes for each compartment (Figure 3B and D). For bacteria, at the phylum level, the phyla identified and relative abundance of each was largely controlled by the compartment as opposed to the rootstock genotype (Figure 3B). For fungi, the pattern was mostly the same, with the greatest differences in taxa and relative abundance of classes being related to compartment (Figure 3D). *Saccharomycetes* showed patterning by rootstock, with all grafted plants showing an increased abundance (Figure 3D). After post-hoc testing we found that berries from the rootstock ‘1103P’ showed significantly higher abundance than ungrafted vines and grafted ‘SO4’ vines (‘1103P’ vs Ungrafted berries: *P* < 0.001 and ‘1103P’ vs ‘SO4’: *P* = 0.043).

### Rootstock and irrigation influence microbiota of winemaking relevance

While characterizing the microbiota of different rootstocks, we observed that samples containing a high relative abundance of *Saccharomycetes* also generally had increased relative abundance of the bacterial order *Acetobacterales*. These taxa, while ubiquitous in and on grape berries, are also causal organisms of the late-season bunch rot disease sour rot and are relevant to winemaking. In order to more thoroughly investigate the impact of rootstock on these agronomically and viticulturally important taxa, we extracted members of the bacterial order *Acetobacterales* and the fungal class *Saccharomycetes* from the rarified dataset. The extracted *Acetobacterales* including 30 species representing seven genera, with most sequence reads comprising members of *Acetobacter* and *Gluconobacter* (6.1% and 83.4%). The extracted *Saccharomycetes* represented 48 species from 12 genera, with most concentrated in the genera *Hanseniaspora* and *Pichia* (32.9% and 25.7%).

Linear models demonstrated that rootstock and its interactions significantly influence the relative abundance of *Acetobacterales* and *Saccharomycetes* (Table 2). *Acetobacterales* relative abundance was significantly influenced by the three way interaction of rootstock, compartment, and irrigation (R×C×I; *P* = 0.002; Table 2). Post-hoc testing showed that all variation was attributable to the berry compartment, with other compartments showing no significant comparisons (Figure 4A). Within the full irrigation treatment, berries of ‘SO4’ grafted vines had elevated relative abundance of *Acetobacterales* in comparison to other vines (SO4 vs Ungrafted *P* = 0.009, ‘SO4’ vs ‘1103P’ *P* = 0.003, and ‘SO4’ vs ‘3309C’ *P* = 0.004). Whereas berries of ‘1103P’ grafted vines had elevated relative abundance under reduced (‘1103P’ vs Ungrafted *P* < 0.001, ‘1103P’ vs ‘3309C’ *P* < 0.001, ‘1103P’ vs ‘SO4’ *P* < 0.001) and no supplemental irrigation treatments (‘1103P’ vs Ungrafted *P* = 0.001, ‘1103P’ vs ‘SO4’ *P < 0*.*001). Saccharomycetes* relative abundance was influenced by the interaction of rootstock and irrigation (R×I; *P* = 0.007; Table 2). Post-hoc testing showed only a single significant comparison for the interaction, ‘1103P’ grafted vines had elevated for *Saccharomycetes* relative abundance compared to ungrafted vines within the reduced irrigation treatment (‘1103P’ vs Ungrafted *P* = 0.048; Figure 4B), however, ‘1103P’ vines compared to ungrafted vines across irrigation treatments showed much higher significance (‘1103P’ vs Ungrafted *P* = 0.006). The relative abundance of *Acetobacterales and Saccharomycetes* for berries were strongly positively correlated (Spearman correlation = 0.735, *P* < 0.001; Figure S6).

**Table 2.**
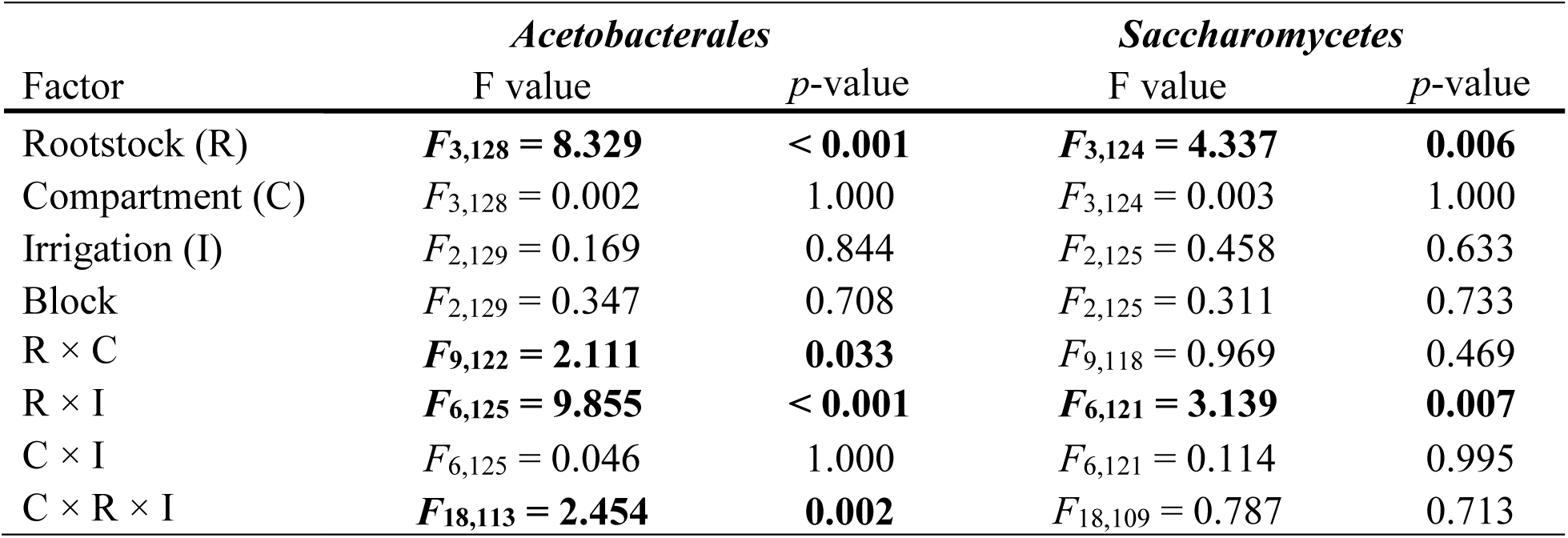
Three-way ANOVA model results for the abundance of *Acetobacterales* and *Saccharomycetes*. Factors that are determined to be significant are bolded and interactions are denoted with the following abbreviations (R) rootstock, (C) compartment, and (I) irrigation.

| Factor | *Acetobacterales* |  | *Saccharomycetes* |  |
| --- | --- | --- | --- | --- |
|  | F value | *p*-value | F value | *p*-value |
| Rootstock (R) | $F_{3,128} = \mathbf{8.329}$ | **< 0.001** | $F_{3,124} = \mathbf{4.337}$ | **0.006** |
| Compartment (C) | $F_{3,128} = 0.002$ | 1.000 | $F_{3,124} = 0.003$ | 1.000 |
| Irrigation (I) | $F_{2,129} = 0.169$ | 0.844 | $F_{2,125} = 0.458$ | 0.633 |
| Block | $F_{2,129} = 0.347$ | 0.708 | $F_{2,125} = 0.311$ | 0.733 |
| R × C | $F_{9,122} = \mathbf{2.111}$ | **0.033** | $F_{9,118} = 0.969$ | 0.469 |
| R × I | $F_{6,125} = \mathbf{9.855}$ | **< 0.001** | $F_{6,121} = \mathbf{3.139}$ | **0.007** |
| C × I | $F_{6,125} = 0.046$ | 1.000 | $F_{6,121} = 0.114$ | 0.995 |
| C × R × I | $F_{18,113} = \mathbf{2.454}$ | **0.002** | $F_{18,109} = 0.787$ | 0.713 |

**Figure 4.**
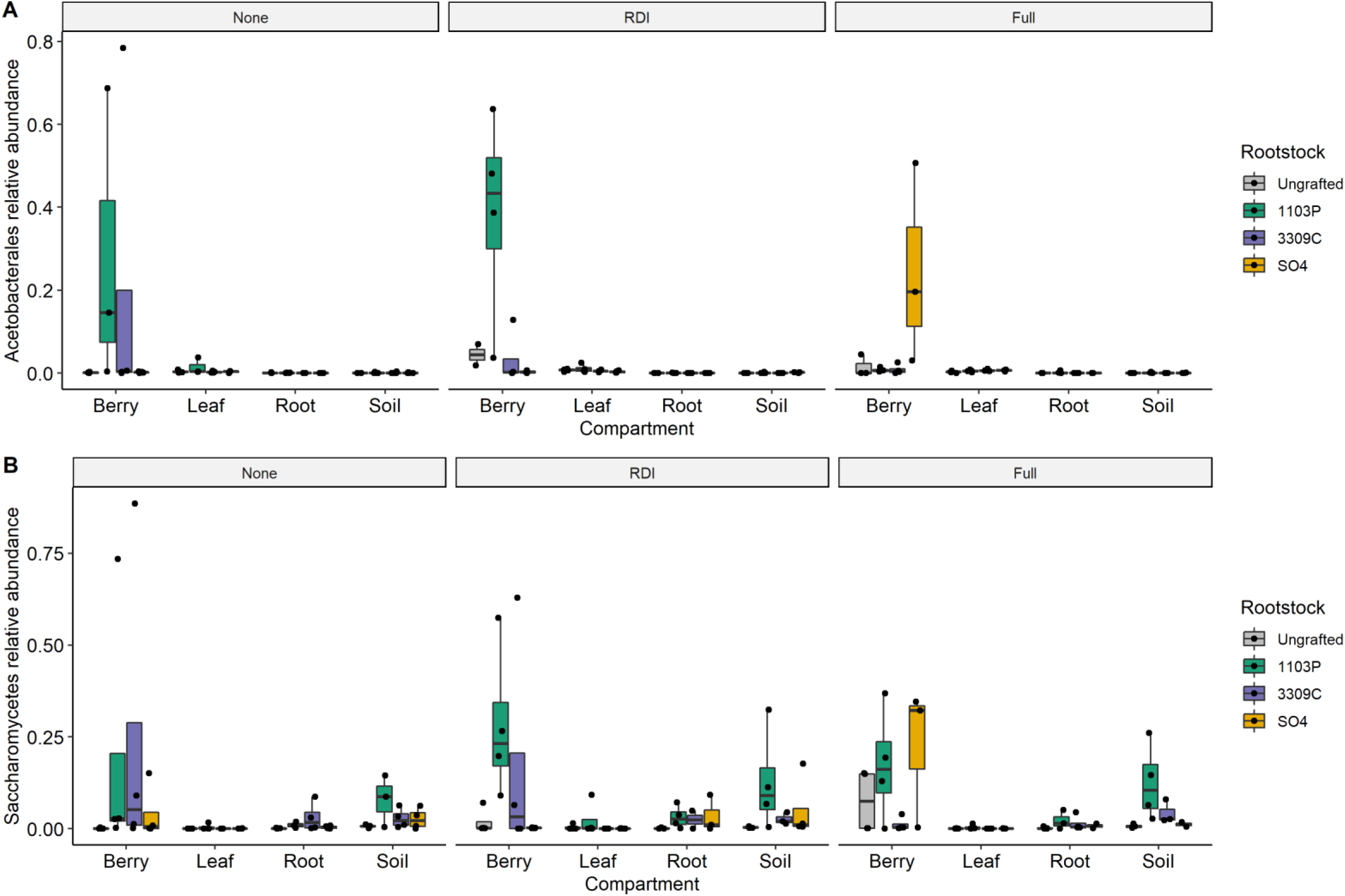
The abundance of A) *Acetobacterales* and B) *Saccharomycetes* are influenced by complex interactions of rootstock genotype, irrigation treatment, and plant compartment. Panels delineate irrigation treatments (None, unirrigated; RDI, reduced replacement of evapotranspiration; Full, Full replacement) and rootstock genotypes correspond to colors in the legend.

### Machine learning and differential abundance analyses

For machine learning-based classification we tested the random forest algorithm on both regions (16S rRNA gene and ITS) both separately and together. We found that the accuracy was similar for each region and the combined dataset, so we report the results of the combined dataset. After hyperparameter tuning of the random forest models (Figure S7, and Table S4) performance on the combined 16s rRNA and ITS testing data showed high accuracy in classifying samples to compartment but not to rootstock (98% and 35% accuracy, respectively; Figure 5A-C). When compartment and rootstock were jointly classified, we found that accuracy was higher (50% accuracy) with most samples being correctly assigned to compartment, but rootstock genotypes assignments were stochastic. Similarly, we were unable to accurately classify which irrigation treatment was applied to a given sample (45% accuracy; Figure S7). These data corroborate other analyses that point to compartment as the primer determinant of microbiome diversity within a vine.

**Figure 5.**
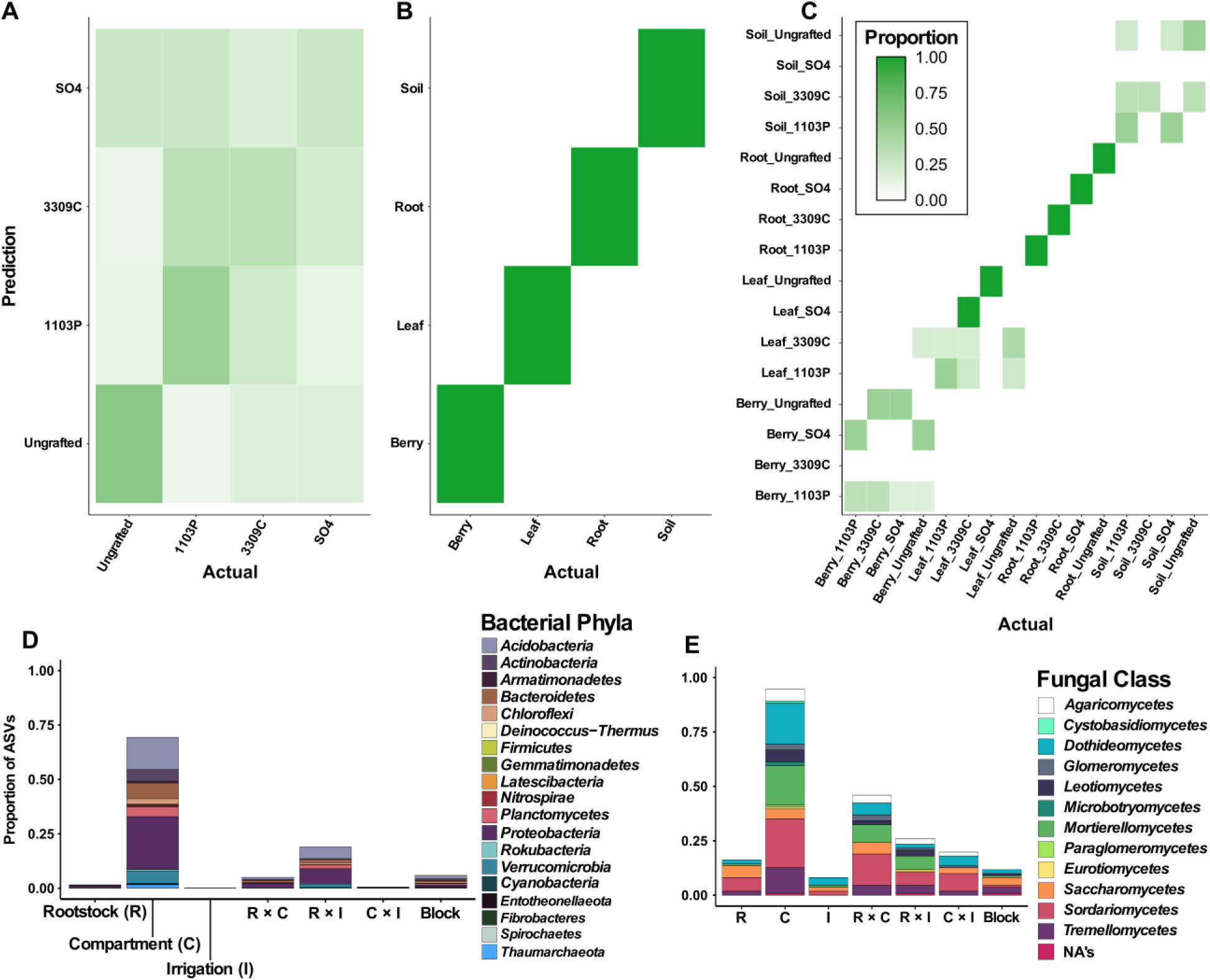
Machine learning was able to distinguish between A & B) the compartment from which the sample was collected but not rootstock genotype to which the vine was grafted and C) when tissue and rootstock were jointly predicted only tissue was able to be predicted accurately. DESeq2 analysis found that multiple sources of variation were responsible for the patterns of differential abundance for amplicon sequence variants (ASVs) of D) bacteria phyla and E) Fungi classes.

Differential abundance analysis in DESeq2 illustrated that ASVs are associated with multiple factors in the experimental design (Figure 5D and E). After filtering ASVs not represented at a depth of least 25 reads in more than 10% of the samples, we were left with 757 bacterial and 111 fungal ASVs. Overall a greater proportion of fungal taxa respond to each of the factors more than bacteria. We found that bacteria and fungi generally followed the same rank order in regards to the proportion of differentially abundant ASVs responding to each source of variation (Figure 5D and E). Tissue showed the largest proportion of differentially abundant ASVs with 69.4% of bacterial and 94.6% of fungal ASVs. For bacteria, interactions between sources of variation showed larger proportions of differentially abundant ASVs than the other main effects (R = 1.5%, I = <1%, R×C = 5.0% and R×I = 19.0%, C×I = <1%; Figure 5D). The main bacterial phyla that showed differential abundance with respect to the experimental design were Proteobacteria, Acidobacteria, Bacteroidetes, Planctomycetes, Actinobacteria, and Chloroflexi (Figure 5D). Fungi also showed a greater proportion of differentially abundant ASVs associated to interaction effects (R = 16.2%, I = 8.1%, R×C = 45.9%, R×I = 19.8%, C×I = 26.1%; Figure 5E). The main fungal classes that responded to the different parts of the experimental design were Dothideomytcetes, Sordariomycetes, Mortierellomycetes, Tremellomycetes, Agaricomycetes, Saccharomycetes, and Leotiomycetes (Figure 5E).

The extracted absolute Log_2_ fold change value of each ASV significantly responding to the experimental design was used to compare the effect sizes of each source of variation. The mean Log_2_ fold change for each of the sources of variation were similar, except for block which had a lower Log_2_ fold change value 1.20 ± 0.20 for bacteria and 1.33 ± 0.28 for fungi (Figure S8). Post-hoc tests showed that for bacteria all comparisons to block were significant. For fungi all comparisons to block were again significant, but we also observed that compartment had a significantly larger Log_2_ fold change mean than rootstock and irrigation (+1.54 and +1.62, respectively).

## Discussion

The grapevine microbiome consists of multiple compartments (soil, root/rhizosphere, leaves, and berries) each of which host unique communities of bacterial and fungal taxa. Within the study, we found that global patterns of alpha and beta diversity indices for both bacteria and fungi, varied primarily by grapevine compartments (Figure 2). Although rootstock genotype exhibits subtle impacts on patterns of bacterial and fungal diversity (Figure 3). Analyses using differential abundance and abundance-based machine learning uncovered additional groups of microorganisms that were associated across the factors and interactions within the experimental design (Figure 5). Notably, rootstock and irrigation interact with rootstock genotype to influence patterns of diversity in specific groups of microorganisms such as *Acetobacterales* and *Saccharomycetes* (Figure 4).

We found that compartments were quite distinct from one another with the abundance of particular bacterial phyla and fungal classes changing dramatically between compartments (Figure 2). In accordance with Zarraonaindia *et al*. (2015), bacterial community structure changed most dramatically between above and belowground samples (Figure 2B). Fungal communities also experienced a strong change between above and belowground samples (Figure 2D) consistent with prior studies (Liu and Howell 2020). The large differences observed between compartments allowed for abundance-based machine learning to easily predict the compartment of a sample (Figure 5B). As we show, the compartments each possess distinct microbiota that might be impacted in diverse ways by the rootstock genotypes.

Soil serves as the main reservoir of microorganisms within agricultural fields (Chi *et al*. 2005; Bulgarelli *et al*. 2013; Zarraonaindia *et al*. 2015; Liu *et al*. 2017; Tkacz *et al*. 2020; Figure 2B and E), as such we expected each of the plant compartments would contain a fraction of the diversity found within the soil. In this study we found that a majority of both the bacterial and fungal ASVs were found in or in association with soil, 74% and 68% respectively (Figure S4). The ASVs not associated with soil could have originated from the atmosphere via rainfall or wind (Ottesen *et al*. 2016) or could be exceedingly rare in the soil so as to avoid detection. Atmospheric microorganism originate from multiple sources, including the local vegetation surfaces (i.e. plant phyllosphere), soils, and bodies of water (Bowers *et al*. 2011; Lymperopoulou *et al*. 2016; Smets *et al*. 2016; Šantl-Temkiv *et al*. 2018). Local atmospheric conditions (e.g. relative humidity, temperature, and etc.) and seasonal variability have been found to influence atmospheric microbial community composition (Fahlgren *et al*. 2010; Bowers *et al*. 2012, 2013; Šantl-Temkiv *et al*. 2018), but strong effects are also associated with the type of land-use (Bowers *et al*. 2011). Hyma and Fay (2013) showed that *Saccharomyces cerevisiae* ecotypes were readily dispersed from oak trees to grapevines in surrounding vineyards. Thus, it is possible some of the microorganisms we identified, only in association with the plant, originated from the surrounding environment and might be important to the formation of aboveground microorganism communities.

The large decrease in diversity observed between below and aboveground compartments can be attributed to multiple factors such as fluctuating water availability, harsh climatic conditions, and limited nutrient access in aboveground compartments (Vorholt 2012). For bacteria and fungi inhabiting aboveground compartments, water availability is paramount to survival and fluctuates greatly with environmental conditions (Beattie 2011; Aung *et al*. 2018). For instance, the level of relative humidity was found to positively correlate with fungal abundance and richness in the air and on leaf surfaces (Talley *et al*. 2002). Harsh climatic conditions, such as exposure to UV radiation (Kadivar and Stapleton 2003), further limit the abundance and richness of microorganisms on the surface of aboveground tissues. In addition, nutrients on the surface of aboveground tissues are scarce and more heterogeneously distributed as compared to nutrients available to soil- and rhizosphere dwelling microorganisms (Leveau and Lindow 2001). These factors have been shown in other studies to limit the bacterial and fungal diversity and likely contribute to the observed diversity levels in both leaf and berry compartment samples in this study.

Our study is the first that attempts to simultaneously assess the root and shoot systems of grapevines with an eye toward understanding the impact of grafting on the shoot system microbiome. Previous research has shown that rootstock genotypes influence the community that associates with the rhizosphere and root endosphere of the grapevine (D’Amico *et al*. 2018; Marasco *et al*. 2018; Berlanas *et al*. 2019). Further, cultivar specific differences in the microbial communities of berries and leaves have been recorded (Bokulich *et al*. 2014, 2016; Mezzasalma *et al*. 2018; Singh *et al*. 2018; Zhang *et al*. 2020). Our results show that grafting and the different rootstock genotypes did not have a large effect on the diversity indices of the different compartments of the vine. Alpha (Figure 3A and C; Table S2 and S3) and beta diversity indices (Table 1) were either non-significant with respect to rootstock and its interactions or only explained a small percentage of variance (1.6% for R and 3.2% for R × I fungal Bray-Curtis; Table 1). This indicates that there is a core microbiome of the sampled grapevines, which is variable by compartment, but is largely conserved across different rootstocks and the process of grafting. This could be the result of multiple factors. First, within our study we did not separate the rhizosphere and root endosphere compartments, opting to grind the root tissue with the rhizosphere still adhered. This could obscure the detection of an effect of rootstock genotype if these compartments, rhizosphere and root endosphere, respond differently. Second, the scion we sampled from, ‘Chambourcin’, is of complex hybrid origin, whereas the previous studies made use of *Vitis vinifera* cultivars (’Barbera’, ‘Tempranillo’, and ‘Lambrusco’; D’Amico *et al*. 2018; Marasco *et al*. 2018; Berlanas *et al*. 2019). Currently, more work is required to understand how the scion portion of a grafted plant influences the microbiota of the rootstock and other compartments of the vine.

While irrigation treatments were applied to the vines throughout the growing season, we did not find a large impact of irrigation on patterns of microbial diversity. There are two different scenarios that can explain these results. The first possibility is that the microbiota associated with the different tissues of the grapevine are unimpacted by the amount of water that the grapevine receives. This is unlikely as previous research has shown that drought typically alters the microbiome across many plant species (Naylor and Coleman-Derr 2018; Fitzpatrick et al. 2018). For example, in a study on Sorghum, under drought conditions rhizosphere and root microbiota showed enrichment of monoderms, bacteria with a single membrane and thick peptidoglycan cell wall, which likely provides increased desiccation resistance (Xu et al. 2018). The second possibility is that the amount of seasonal precipitation received during the growing season was enough to obscure some of the signal from the irrigation treatment. Previous research in this experimental vineyard from prior years, have shown that irrigation treatments have impacted multiple phenotypes, including, physiologic measurements in 2014-2015 (Maimaitiyiming *et al*. 2017) and the ionome and morphology of leaves in 2014-2016 (Migicovsky *et al*. 2019). However, the effects of irrigation in the past in this vineyard were weaker than other parts of the experimental design (e.g. development and rootstock genotype) and these studies collected measurements or samples across a wider portion of the growing season than the current study. In the month prior to the sampling date (Aug. 18 - Sept. 18, 2018) the vineyard experienced an average precipitation of 0.56 ± 1.27 cm of precipitation per day with three days that experienced greater than 3 cm of precipitation (Missouri Historical Agricultural Weather Database URL: http://agebb.missouri.edu/weather/history/). This amount of precipitation exceeded the amount of water the soil was expected to lose from evapotranspiration, given the environmental conditions, leading to vines not experiencing severe water stress.

Sour rot is a disease linked to the four way interaction of a susceptible grapevine host, *Drosophila* spp., acetic acid bacteria (e.g. *Acetobacter* spp. and *Gluconobacter* spp.) and various yeast species (*Hanseniaspora uvarum, Candida* spp., and *Pichia* spp.; Hall *et al*. 2018). This disease is of particular importance in growing regions that experience high rainfall during late stages of berry maturation such as the middle USA. Current strategies to combat the disease recommend use of insecticides in order to control the population of *Drosophila* and physical measures to prevent damage to berry clusters (e.g. bird netting; Wilcox *et al*. 2015; Hall, Loeb, and Wilcox 2018; Sun *et al*. 2019). Until recently, *Drosophila* was thought to contribute to the pathogenesis of sour rot by serving as the vector of the above described microorganisms and delaying wound healing of damaged berries (Barata, Pais, *et al*. 2011; André Barata *et al*. 2012). However, Hall et al. (2018a and 2018b) showed that sour rot symptoms develop in the presence of both wild-type and axenic (germ-free) *Drosophila*, but not in exclusion of *Drosophila*, and that acetic acid bacteria and yeast species can be readily cultured from asymptomatic berry clusters. Given that the microorganisms implicated in sour rot originate from the grape clusters themselves, we investigated whether particular rootstocks contributed to the enrichment of these groups of microorganisms, *Acetobacterales* and *Saccharomycetes*.

We found both *Acetobacterales* and *Saccharomycetes* were influenced by complex interactions involving multiple factors including rootstock, irrigation, and compartment (Figure 4). For *Acetobacterales*, we found a significant three-way interaction between rootstock, irrigation, and compartment (Table 2). This was driven by an elevated relative abundance in berries for vines grafted on the rootstock ‘1103P’ in the unirrigated and reduced irrigation treatments and by ‘SO4’-grafted vines in the full irrigation treatment (Figure 4A). For *Saccharomycetes*, we found a significant interaction between rootstock and irrigation but post-hoc testing showed only a single significant comparison whereas the main effect of rootstock more significant post-hoc comparisons and explained a similar amount of the variance (7.69% and 7.07%, respectively; Table 2). *Saccharomycetes* across irrigation treatments showed elevated relative abundances in grafted vines compared to ungrafted (Figure 4B). We also observed a strong positive correlation between the relative abundance of *Acetobacterales* and *Saccharomycetes* for the berry compartment (Figure S6).

This correlation indicates that these groups likely have mutualistic interactions. Previous pathology work in sour rot, has found that ethanol levels in berries rise prior to accumulation of acetic acid (André Barata *et al*. 2012; Hall, Loeb, Cadle-Davidson, *et al*. 2018). The hypothesized mechanism involves yeast, first fermenting sugars within the berries producing ethanol as a by-product, which then undergoes oxidative fermentation by acetic acid bacteria to produce acetic acid. This two-step process might explain why we observed *Saccharomycetes* across more samples than *Acetobacterales*, as *Saccharomycetes* might be a prerequisite for accumulation of *Acetobacterales* by producing a microenvironment within the berry more suitable for growth.

Both of the microorganism groups tended to be in higher abundance in the berry compartment of grafted grapevines. This indicates that either grafting or the rootstock genotypes in this study have the ability to influence the abundance of the microorganisms implicated in sour rot. Previous work has shown that plant genotypes possess unique root exudate profiles (Micallef *et al*. 2009; Lundberg *et al*. 2012; Mönchgesang *et al*. 2016) and that these exudates shape microbial communities (Sasse *et al*. 2018; Zhalnina *et al*. 2018). Thus, metabolomic analysis would be useful to corroborate the association that we observed with ‘1103P’ and the grafted vines in comparison to ungrafted vines. It is possible that rootstocks produce compounds, whether specific to genotype or as an effect of grafting, that contribute to elevated abundance of *Acetobacterales* and *Saccharomycetes* either directly through specialized substrates or indirectly via suppression of competition with other microorganisms. With the recent emergence of insecticide resistance in populations of *Drosophila* reported in the Finger Lakes region, New York (Sun *et al*. 2019), additional understanding of the dynamics of the constituents of sour rot to common vinicultural techniques are a critical need.

These microorganisms also contribute to wine quality. The acetic acid bacteria we recovered were mostly of the genera *Gluconobacter* and *Acetobacter*. These are aerobic bacteria able to oxidize sugars to acetic acid when berries are damaged (Hall, Loeb, Cadle-Davidson, *et al*. 2018) and during wine production when products are exposed to oxygen (Bartowsky *et al*. 2003; Bartowsky and Henschke 2008; Barata, Campo, *et al*. 2011). Non-*Saccharomyces* yeasts have been investigated for the properties in winemaking (Ciani and Maccarelli 1997; Jolly *et al*. 2014; Padilla *et al*. 2016; Ciani and Comitini 2019). The non-*Saccharomyces* yeast we recovered were mostly *Hanseniaspora* and *Pichia*, both of which are generally considered spoilage yeasts when they are dominate within a fermentation (A. Barata *et al*. 2012) but can be beneficial if allowed to perform initial fermentation and then followed up with an inoculation of a strong fermenting yeast species (e.g. *Saccharomycetes cerevisiae*; Jolly *et al*. 2014). *Hanseniaspora* in fermentations was found to contribute to the accumulation of acetic acid and ethyl acetate (Viana *et al*. 2008; Padilla *et al*. 2016; Ciani and Comitini 2019). Similarly, *Pichia* can contribute to higher levels of ethyl acetate in wines (Viana *et al*. 2008) although a study using simple culture mediums show conflicting results (Moreira *et al*. 2005). These genera have also been used to successfully enhance wine fermentations when co-inoculated with another fermenting yeast, due to their β–glucosidase activity allowing for higher productions of volatiles (Rodriguez *et al*. 2004; Hernández-Orte *et al*. 2008; Swangkeaw *et al*. 2011). Thus, our results illustrate that the treatments imposed on the vines in the vineyard, namely grafting and irrigation, can lead to microbial changes in the berries which could have further implications on the fermentation microbiome.

## Conclusion and future directions

Grafting, the process by which plant parts are fused together, provides a unique avenue to explore the ways in which root and shoot systems interact. In grapevines, we have shown that plant compartments retain unique bacterial and fungal communities regardless of the whether they were grafted or to which rootstock genotype they are grafted. Indicating that the environmental conditions that microorganisms are exposed to within different parts of the plant are paramount. We found that rootstock genotype, irrigation, and their interaction had small effects on global patterns but showed significant associations with groups of microorganisms such as *Acetobacterales* and *Saccharomycetes*, which are implicated in the disease sour rot and can contribute to wine characteristics within the fermentation process. This result will require further experimental validation in order to understand whether the associations with these microorganisms impact the susceptibility of clusters on an individual vine to sour rot and how the characteristics of the resulting wines are impacted. In addition, the role of the scion of grafted grapevines in shaping a vine’s microbial communities warrants investigation. An experimental design that makes use of reciprocal scion and rootstock combinations would allow for isolating the roles of the root and shoot systems in regulating the formation of a stable microbiome.

## Supplementary information

Additional file 1:

**Figure S1**. Taxonomic barplots for negative and positive controls.

**Figure S2**. Rarefaction curves for bacterial and fungal samples.

**Figure S3**. Alpha diversity metrics for bacterial and fungal samples

**Figure S4**. Venn diagrams show the overlap of ASVs between compartments for bacterial and fungal samples

**Figure S5**. Principal coordinates analysis showing the third axis based on bacterial unweighted Unifrac distance and fungal Bray-Curtis distance.

**Figure S6**. Scatterplot of the abundance of *Acetobacterales* and *Saccharomycetes*, showing a strong positive correlation.

**Figure S7**. Out of bag error estimate across values of the number of trees models attempting to predict rootstock, compartment, and rootstock by compartment jointly.

**Figure S8**. Absolute Log2 fold change values for ASVs associated with each source of variation.

**Figure S9**. Confusion matrix of model attempting to predict irrigation treatment.

**Table S1**. Soil chemical analysis for the experimental vineyard.

**Table S2**. Anova tables for bacterial alpha diversity metricss

**Table S3**. Anova tables for fungal alpha diversity metrics

**Table S4**. Model hyperparameters picked as optimal after tuning in Caret.

**Table S5**. Output statistics for machine learning model predicting rootstock genotype.

**Table S6**. Output statistics for machine learning model predicting compartment.

**Table S7**. Output statistics for machine learning model jointly predicting rootstock and compartment.

## Availability of data and code

All raw sequencing data is available on NCBI under BioProject ID PRJNA647904 and SRA accessions SRR12304683-SRR12304870. All code to reproduce the analysis and figures is available on GitHub at https://github.com/Kenizzer/Grapevine_Microbiota.

## Author Contributions

JFS, MEH, MTK, and AJM designed the experimental and collection design. JFS, MTK, ZNH, and AJM performed sample collection. JFS performed sample processing and data analysis. All authors contributed to data interpretation, the writing of the manuscript, and approved the final draft.

## Acknowledgements

This material is based upon work supported by the National Science Foundation Graduate Research Fellowship to JFS under Grant No. 1758713 and NSF 1546869 to AJM. This research was also funded by a grant from the Missouri Grape and Wine Institute to AJM, MEH, MTK, and JFS. The authors would like to thank Courtney Coleman, Emma Frawley, Laura Klein, and Laszlo Kovacs for assistance in sample collection and members of the Miller lab for helpful comments in preparing this article.

