## Supplemental Materials file 1 for "Grapevine microbiota reflect diversity among compartments and complex interactions within and among root and shoot systems"

**Figure S1.** Taxonomic barplots for negative and positive controls used within the study. Above each bar is the number of reads after quality filtering for each of the samples. Mock DNA samples were ZymoBIOMICS™ Microbial community DNA standard (Cat No. D6305). Samples without a bar represent samples for which no reads passed quality filtering.

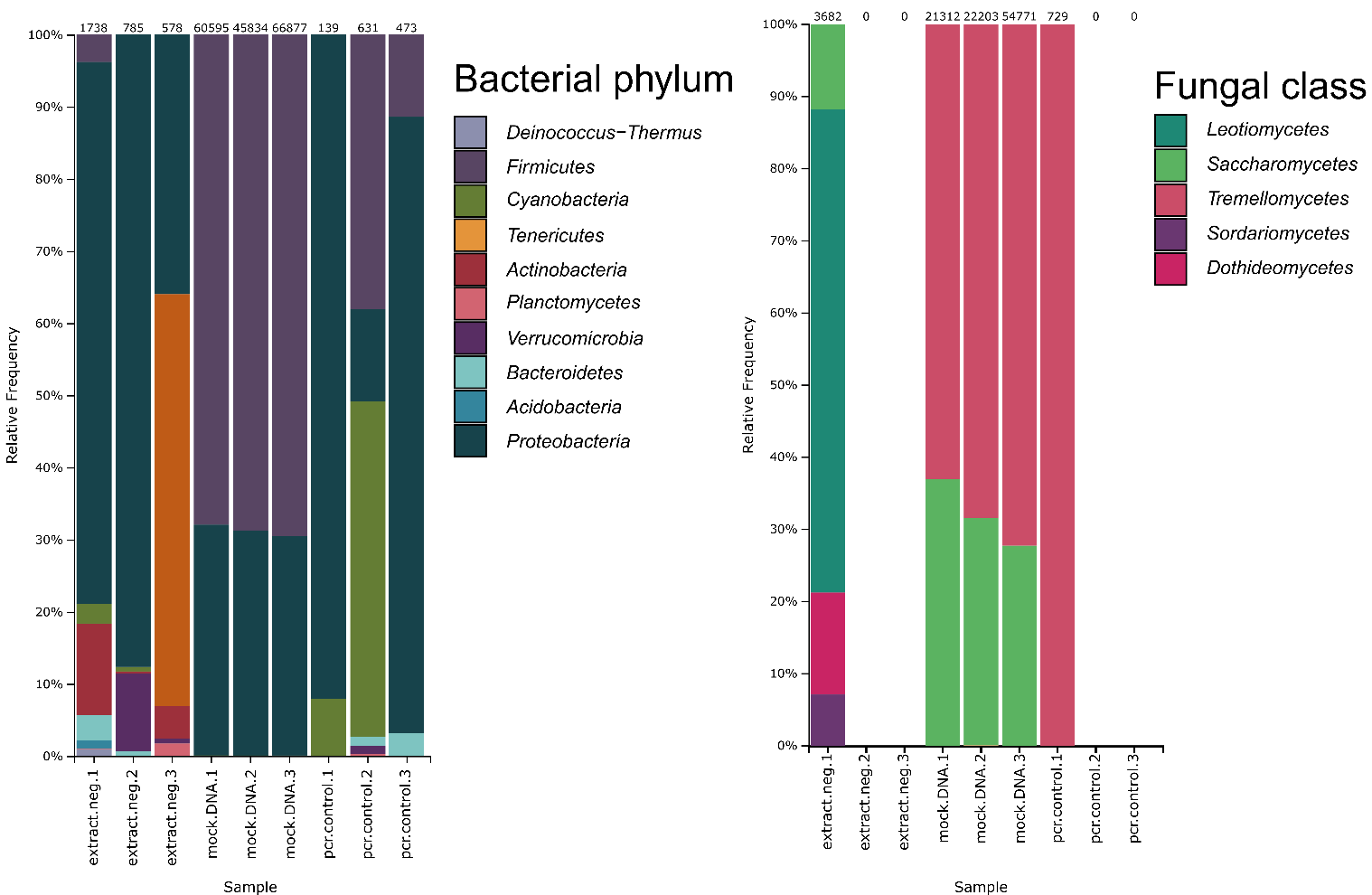

**Figure S2.** Rarefaction curves for A) bacterial and B) fungal samples.

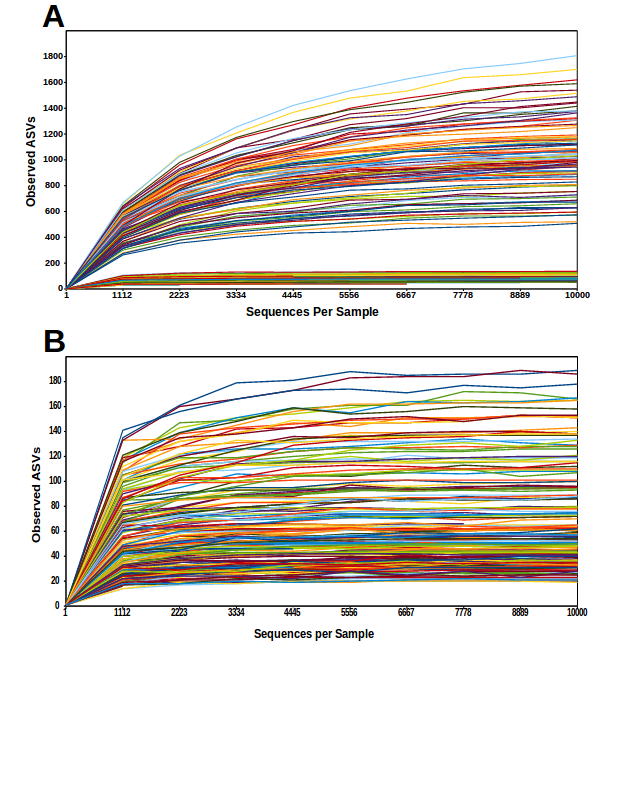

**Figure S3.** Alpha diversity metrics for A-C) bacterial and D-E) fungal communities by compartment.

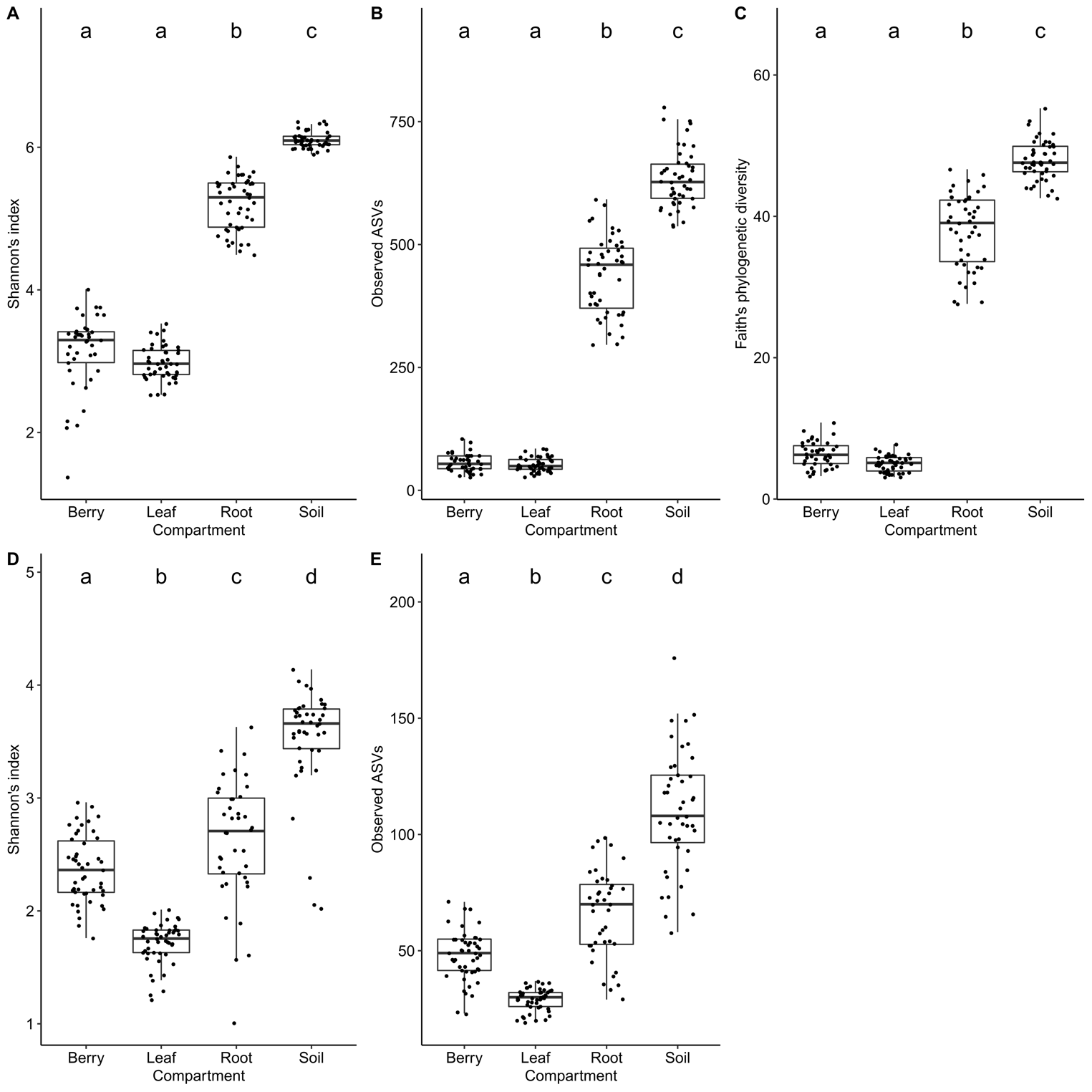

**Figure S4.** Venn diagrams depicting the overlap of ASVs between compartments and soil for A) bacterial and B) fungal samples.

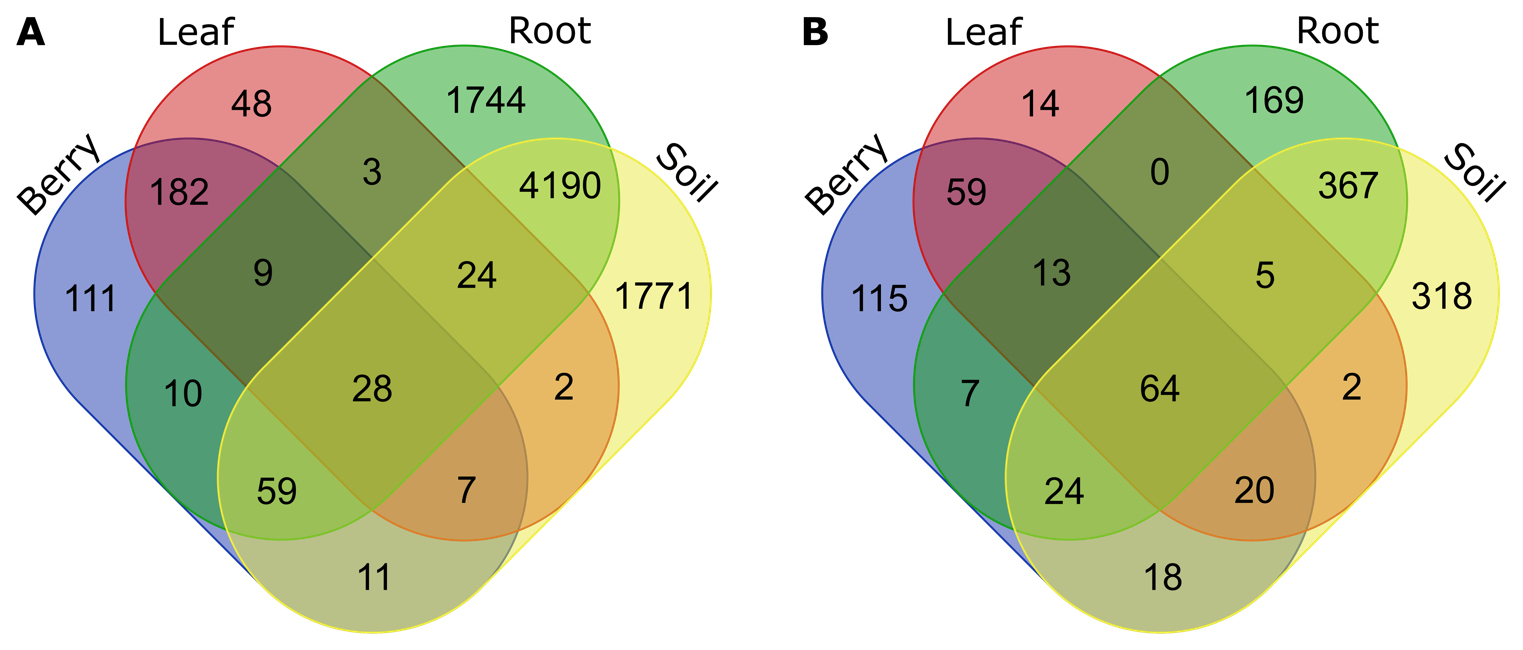

**Figure S5.** Principal coordinates analysis for A & B) Bacterial communities based on unweighted UniFrac distance and C & D) fungal communities based on Bray-Curtis distance.

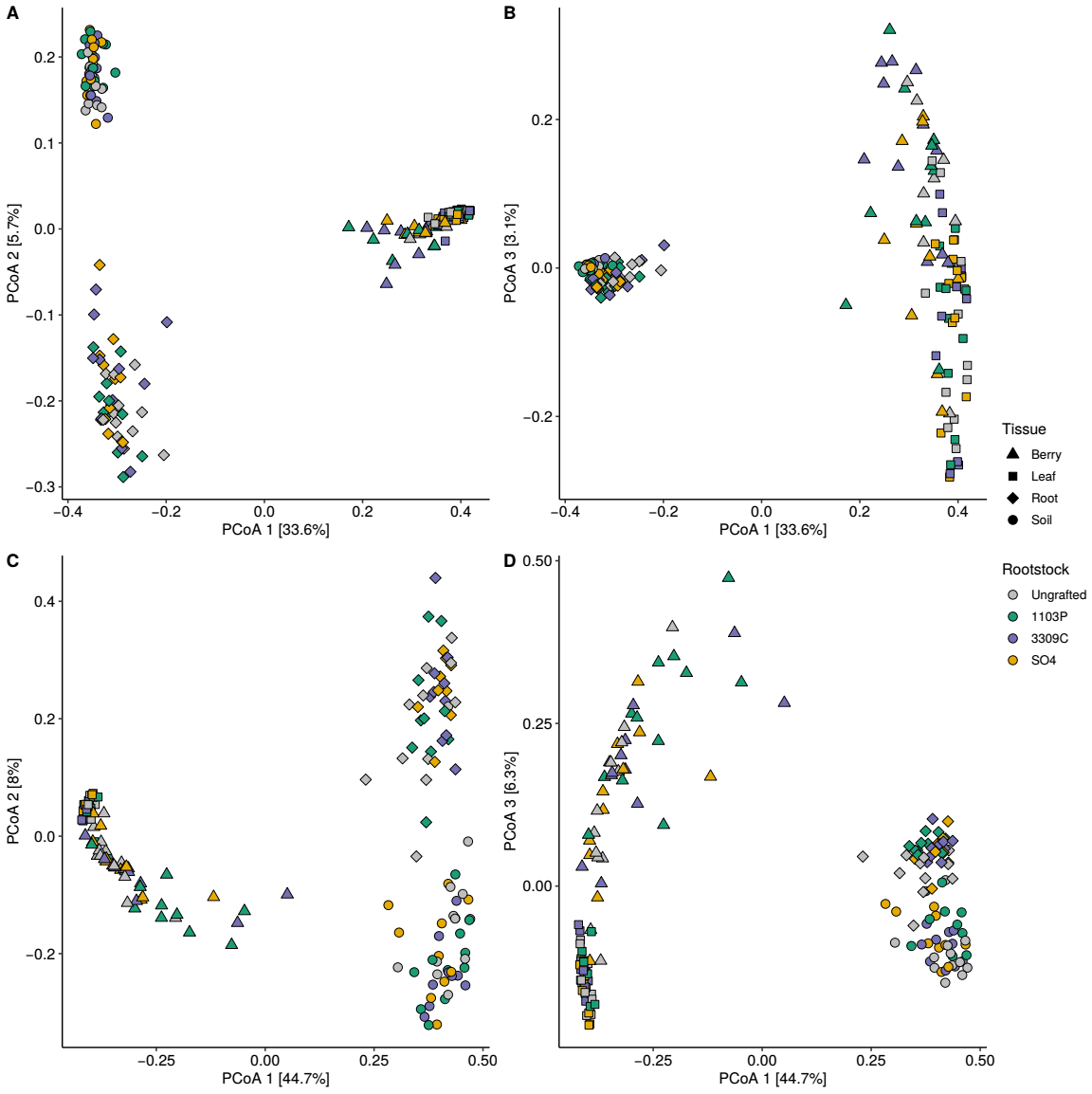

**Figure S6.** The abundance of *Acetobacterales* and *Sacchromycetes* show a strong positive correlation in the berry compartment. Spearman correlation coefficient and *P* value are reported for each compartment.

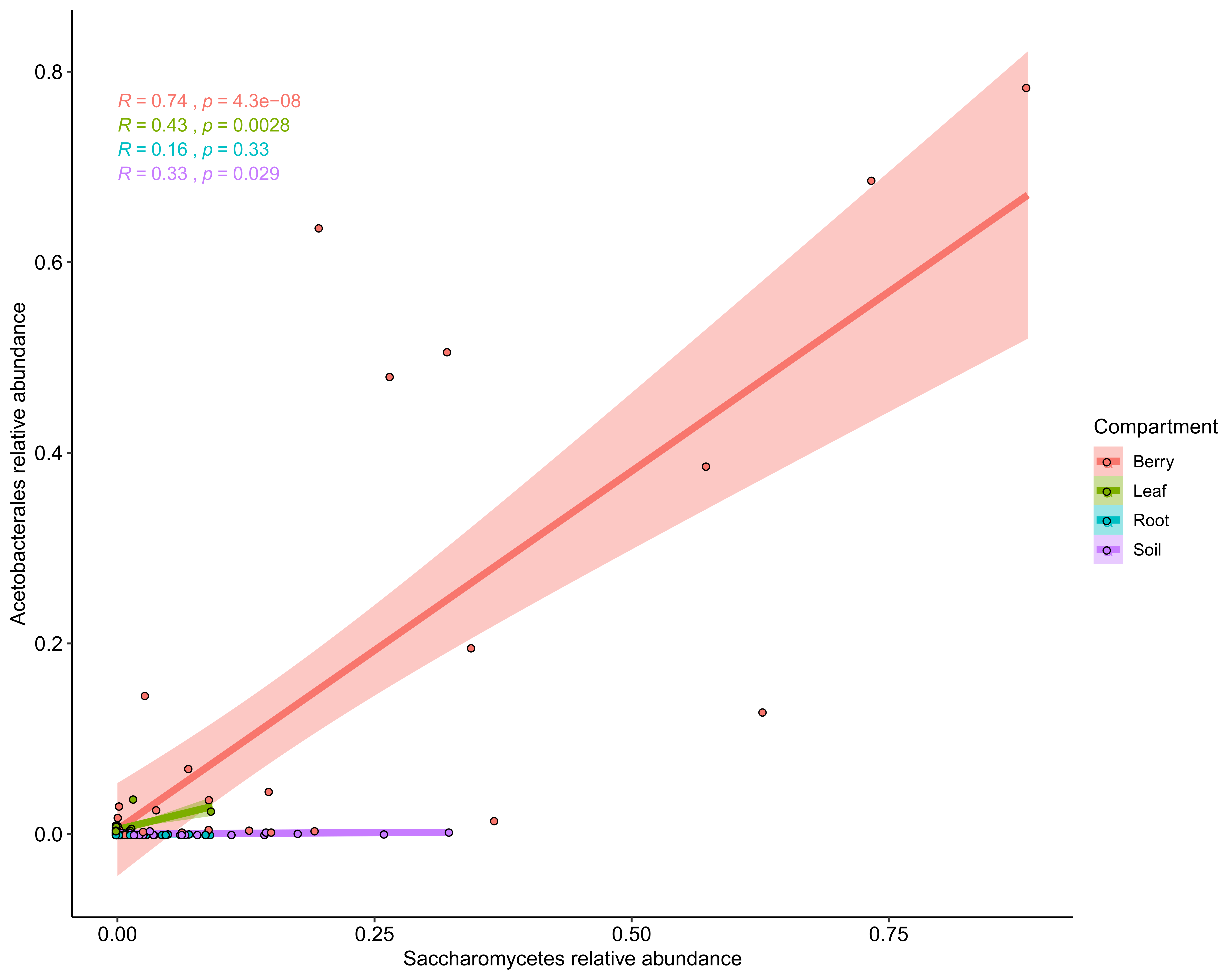

**Figure S7.** Out of Bag error estimate for different values of the number of trees across models attempting to predict rootstock, compartment, and rootstock by compartment jointly.

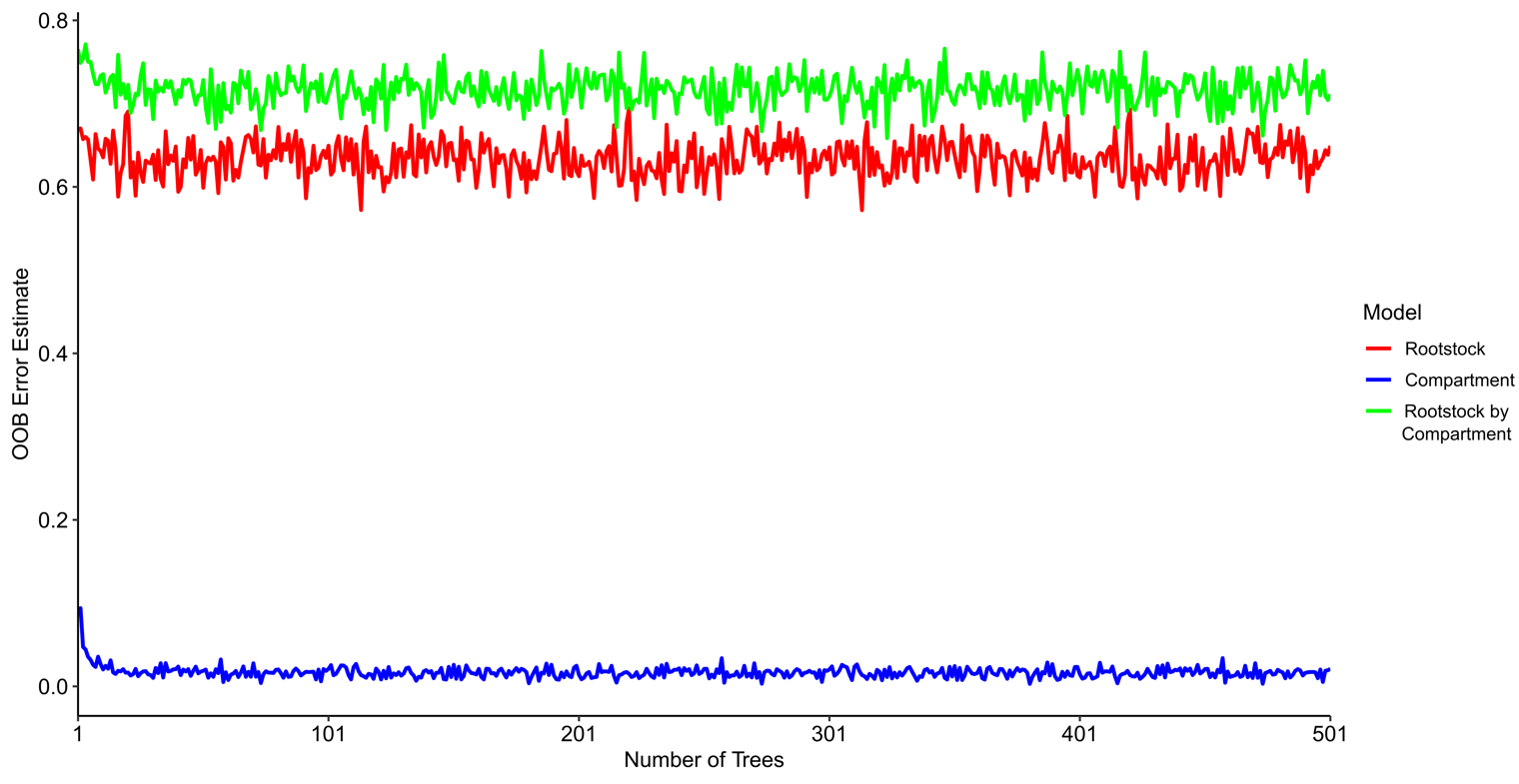

**Figure S8.** Violin plot of extracted absolute Log_2_ fold change values for ASVs associated with each source of variation.

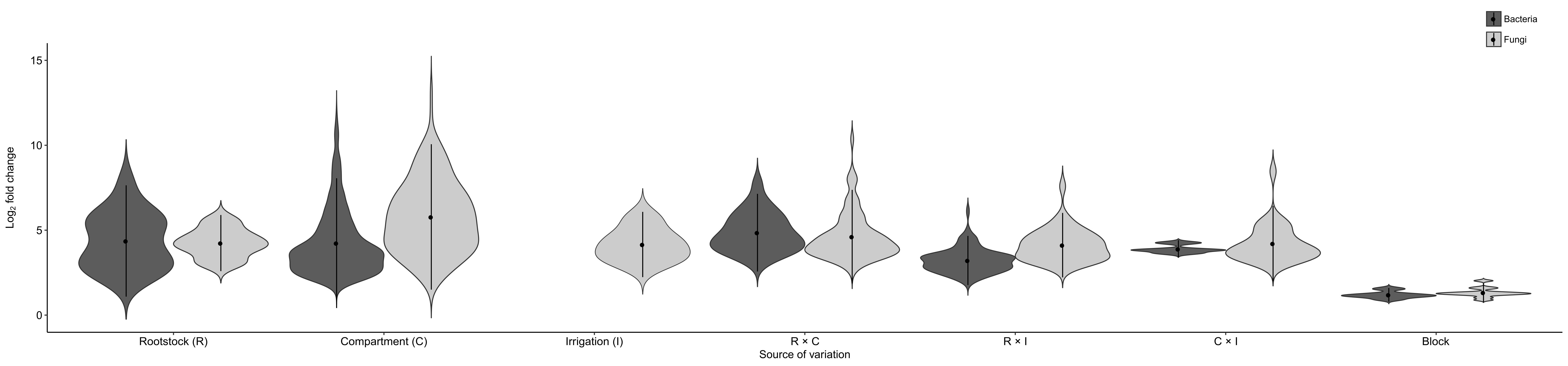

**Figure S9.** Machine learning analysis attempting to predict the irrigation treatment a given sample was under. Irrigation treatments were full replacement of evapotranspiration, intermediate replacement (50%), and nonirrigated.

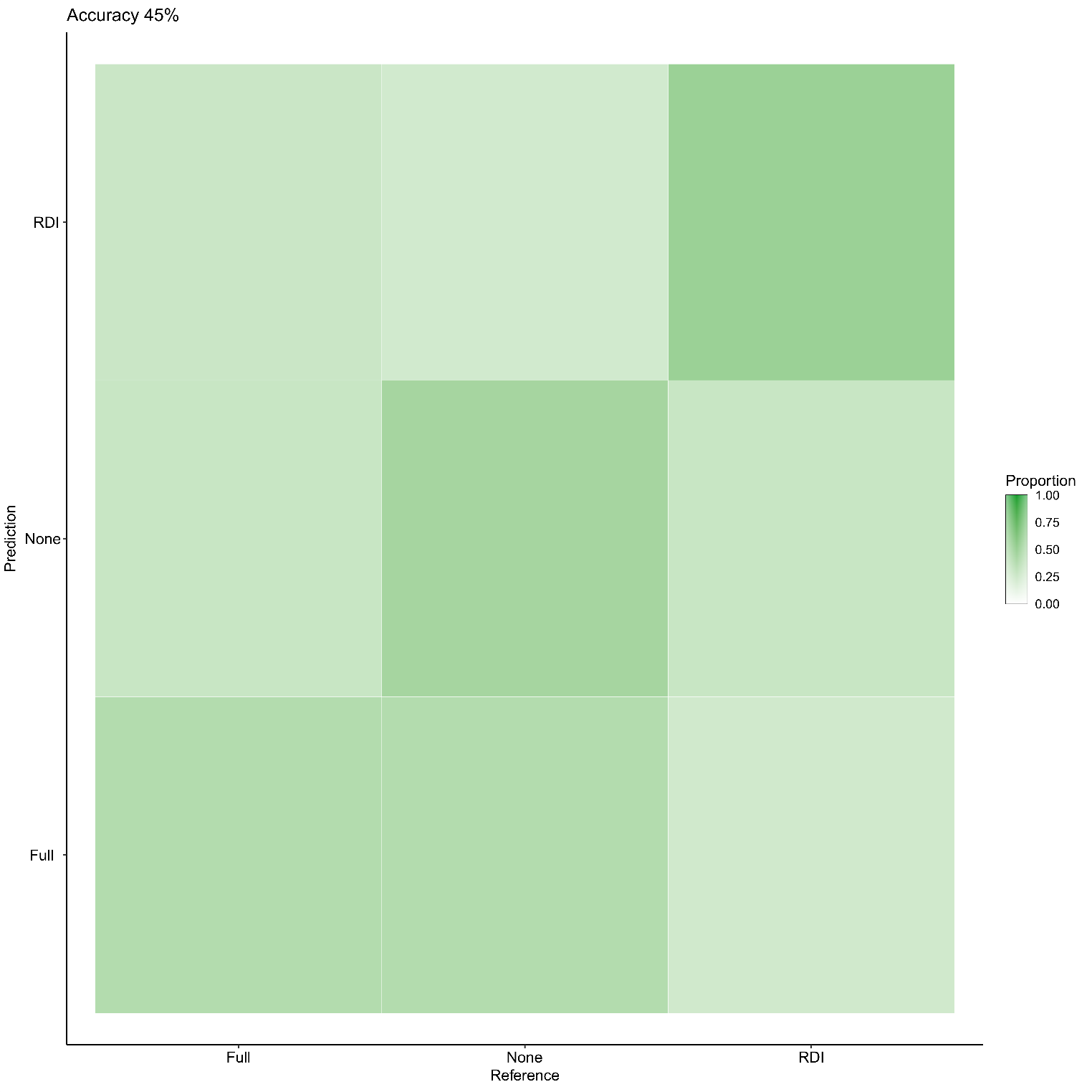

**Table S1.** Soil analysis of samples collected from the University of Missouri Southwest Research Center in Mount Vernon, MO. Analysis was completed at the University of Arkansas Fayetteville Agricultural Diagnostic Laboratory according to the established protocols (Sikora and Kissel 2014; Zhang et al. 2014). Values are reported in mg/kg (ppm) with the exception of pH which is reported in the standard scale.

| **Sample** | **pH** | **P** | **K** | **Ca** | **Mg** | **S** | **Na** | **Fe** | **Mn** | **Zn** | **Cu** | **B** |
| --- | --- | --- | --- | --- | --- | --- | --- | --- | --- | --- | --- | --- |
| **Block 3N** | 7.2 | 123 | 197 | 2590 | 311 | 12 | 7 | 82 | 83 | 60 | 3.4 | 1.5 |
| **Block 3I** | 7.4 | 128 | 130 | 2873 | 296 | 11 | 14 | 83 | 76 | 30 | 4.4 | 1.2 |
| **Block 3F** | 7.0 | 138 | 131 | 2633 | 287 | 13 | 12 | 99 | 65 | 36 | 5.5 | 1.5 |
| **Block 2I** | 7.1 | 112 | 124 | 2793 | 297 | 14 | 15 | 82 | 99 | 49 | 33.2 | 1.7 |
| **Block 2F** | 7.2 | 95 | 100 | 2502 | 321 | 11 | 15 | 93 | 62 | 35 | 7.5 | 1.4 |
| **Block 2R** | 7.2 | 189 | 191 | 2849 | 314 | 14 | 7 | 104 | 76 | 34 | 15.1 | 1.6 |
| **Block 1I** | 7.5 | 178 | 127 | 3302 | 311 | 12 | 16 | 90 | 83 | 50 | 11.2 | 1.8 |
| **Block 1N** | 7.1 | 178 | 160 | 2798 | 262 | 12 | 7 | 110 | 69 | 27 | 10.3 | 1.5 |
| **Block 1F** | 7.3 | 186 | 154 | 2857 | 348 | 14 | 14 | 104 | 81 | 30 | 12.3 | 1.6 |

**Table S2.** Anova tables for bacterial alpha diversity metrics.

| **Faith's phylogenetic diversity** | | | | **Inverse Simpson's index** | | |
| --- | --- | --- | --- | --- | --- | --- |
| Factor | SS | *F* value | *p*-value | SS | *F* value | *p*-value |
| Intercept | **96560.477** | ***F*_1,180_ = 9936.466** | **< 0.001** | **1493749.18** | ***F*_1,180_ = 1607.342** | **< 0.001** |
| Compartment (C) | **64538.239** | ***F*_3,178_ = 2213.749** | **< 0.001** | **2357695.10** | ***F*_3,178_ = 845.662** | **< 0.001** |
| Rootstock (R) | 32.471 | *F*_3,178_ = 1.114 | 0.346 | 3528.06 | *F*_3,178_ = 1.265 | 0.289 |
| Irrigation (I) | 47.609 | *F*_2,179_ = 2.450 | 0.090 | 1625.56 | *F*_2,179_ = 0.875 | 0.419 |
| Block | **69.621** | ***F*_2,179_ = 3.582** | **0.031** | **6220.15** | ***F*_2,179_ = 3.347** | **0.038** |
| C X R | 72.984 | *F*_9,172_ = 0.834 | 0.586 | 4075.24 | *F*_9,172_ = 0.487 | 0.881 |
| C X I | 122.987 | *F*_6,175_ = 2.109 | 0.056 | 4011.82 | *F*_6,175_ = 0.719 | 0.635 |
| R X I | 73.418 | *F*_6,175_ = 1.259 | 0.281 | 3806.05 | *F*_6,175_ = 0.683 | 0.664 |
| C X R X I | 89.634 | *F*_18,163_ = 0.5124 | 0.948 | 14491.03 | *F*_18,163_ = 0.866 | 0.620 |
| Residuals | 1273.030 |  |  | 121742.09 |  |  |

| **Observed ASVs** | | | | **Shannon's Diversity** | | |
| --- | --- | --- | --- | --- | --- | --- |
| Factor | SS | *F* value | *p*-value | SS | *F* value | *p*-value |
| Intercept | **14178127.06** | ***F*_1,180_ = 5311.543** | **< 0.001** | **3109.42** | ***F*_1,180_ = 24224.460** | **< 0.001** |
| Compartment (C) | **11212803.39** | ***F*_3,178_ = 1400.215** | **< 0.001** | **321.84** | ***F*_3,178_ = 835.780** | **< 0.001** |
| Rootstock (R) | 6316.28 | *F*_3,178_ = 0.789 | 0.502 | 0.08 | *F*_3,178_ = 0.195 | 0.899 |
| Irrigation (I) | 7320.21 | *F*_2,179_ = 1.371 | 0.257 | 0.04 | *F*_2,179_ = 0.148 | 0.862 |
| Block | 13075.72 | *F*_2,179_ = 2.449 | 0.090 | 0.28 | *F*_2,179_ = 1.075 | 0.344 |
| C X R | 12698.48 | *F*_9,172_ = 0.529 | 0.852 | 0.24 | *F*_9,172_ = 0.206 | 0.993 |
| C X I | 17678.90 | *F*_6,175_ = 1.104 | 0.364 | 0.64 | *F*_6,175_ = 0.833 | 0.546 |
| R X I | 22167.44 | *F*_6,175_ = 1.384 | 0.226 | 1.04 | *F*_6,175_ = 1.355 | 0.238 |
| C X R X I | 31377.77 | *F*_18,163_ = 0.653 | 0.851 | 1.52 | *F*_18,163_ = 0.658 | 0.846 |
| Residuals | 349678.95 |  |  | 16.81 |  |  |

**Table S3.** Anova tables for fungal alpha diversity metrics.

| **Inverse Simpson's index** | | | | **Observed ASVs** | | | **Shannon's Diversity** | | |
| --- | --- | --- | --- | --- | --- | --- | --- | --- | --- |
| Factor | SS | *F* value | *p*-value | SS | *F* value | *p*-value | SS | *F* value | *p*-value |
| Intercept | **13074.903** | ***F*_1,176_ = 602.832** | **<0.001** | **625159.756** | ***F*_1,176_ = 2236.156** | **<0.001** | **1010.141** | ***F*_1,176_ = 6451.465** | **<0.001** |
| Compartment (C) | **6559.548** | ***F*_3,174_ = 100.812** | **<0.001** | **161851.084** | ***F*_3,174_ = 192.977** | **<0.001** | **74.877** | ***F*_3,174_ = 159.407** | **0.001** |
| Rootstock (R) | **219.702** | ***F*_3,174_ = 3.377** | **0.020** | 1454.586 | *F*_3,174_ = 1.734 | 0.163 | 0.527 | *F*_3,174_ = 1.122 | 0.343 |
| Irrigation (I) | 87.568 | *F*_2,175_ = 2.019 | 0.137 | 1288.001 | *F*_2,175_ = 2.304 | 0.104 | 0.661 | *F*_2,175_ = 2.112 | 0.125 |
| Block | 115.236 | *F*_2,175_ = 2.657 | 0.074 | 555.667 | *F*_2,175_ = 0.994 | 0.373 | 0.697 | *F*_2,175_ = 2.224 | 0.112 |
| C X R | **427.601** | ***F*_9,168_ = 2.191** | **0.027** | 3466.575 | *F*_9,168_ = 1.378 | 0.205 | 0.846 | *F*_9,168_ = 0.600 | 0.795 |
| C X I | 96.701 | *F*_6,171_ = 0.743 | 0.616 | 2182.995 | *F*_6,171_ = 1.301 | 0.261 | 1.129 | *F*_6,171_ = 1.202 | 0.310 |
| R X I | 38.709 | *F*_6,171_ = 0.297 | 0.937 | 1668.859 | *F*_6,171_ = 0.995 | 0.432 | 0.625 | *F*_6,171_ = 0.666 | 0.678 |
| C X R X I | 212.811 | *F*_18,159_ = 0.545 | 0.931 | 4385.173 | *F*_18,159_ = 0.871 | 0.613 | 1.740 | *F*_18,159_ = 0.617 | 0.881 |
| Residuals | 2754.520 |  |  | 35505.249 |  |  | 19.885 |  |  |

**Table S4.** Selected model hyperparameters showing optimal results in Caret.

| **Model** | **mtry** | **Minimum node size** | **Number of trees** |
| --- | --- | --- | --- |
| **Rootstock** | 2827 | 5 | 314 |
| **Compartment** | 949 | 5 | 381 |
| **Rootstock × Compartment** | 7522 | 10 | 324 |

**Table S5.** Output statistics for machine learning model predicting rootstock genotype.

| **Prediction: Rootstock** | | | | |
| --- | --- | --- | --- | --- |
| **Class** | **Ungrafted** | **1103P** | **3309C** | **SO4** |
| **Precision** | 0.583 | 0.500 | 0.333 | 0.273 |
| **Recall** | 0.583 | 0.364 | 0.333 | 0.375 |
| **F1** | 0.583 | 0.421 | 0.333 | 0.316 |

**Table S6.** Output statistics for machine learning model predicting compartment.

| **Prediction: Compartment** | | | | |
| --- | --- | --- | --- | --- |
| **Class** | **Berry** | **Leaf** | **Root** | **Soil** |
| **Precision** | 1 | 1 | 1 | 1 |
| **Recall** | 1 | 1 | 1 | 1 |
| **F1** | 1 | 1 | 1 | 1 |

**Table S7.** Output statistics for machine learning model jointly predicting rootstock and compartment.

| **Prediction: Rootstock by Compartment** | | | | | | | | | | | | | | | | |
| --- | --- | --- | --- | --- | --- | --- | --- | --- | --- | --- | --- | --- | --- | --- | --- | --- |
| **Class** | **Berry 1103P** | **Berry 3309C** | **Berry SO4** | **Berry Ungrafted** | **Leaf 1103P** | **Leaf 3309C** | **Leaf SO4** | **Leaf Ungrafted** | **Root 1103P** | **Root 3309C** | **Root SO4** | **Root Ungrafted** | **Soil 1103P** | **Soil 3309C** | **Soil SO4** | **Soil Ungrafted** |
| **Precision** | 0.333 | NA | 0 | 0 | 0.5 | 0.200 | 0 | 0 | 1 | 1 | 1 | 1 | 0.500 | 0.333 | NA | 0.500 |
| **Recall** | 0.667 | 0 | 0 | 0 | 0.667 | 0.333 | 0 | 0 | 1 | 1 | 1 | 1 | 0.333 | 1 | 0 | 0.667 |
| **F1** | 0.444 | NA | NA | NA | 0.571 | 0.250 | NA | NA | 1 | 1 | 1 | 1 | 0.400 | 0.500 | NA | 0.571 |
